# Alternatives to photorespiration: A systems-level analysis reveals mechanisms of enhanced plant productivity

**DOI:** 10.1101/2024.10.16.618214

**Authors:** Edward N. Smith, Marvin van Aalst, Andreas P.M. Weber, Oliver Ebenhöh, Matthias Heinemann

## Abstract

Photorespiration causes a significant decrease in crop yield due to mitochondrial decarboxylation. Alternative pathways (APs) have been designed to relocate the decarboxylating step or even fix additional carbon. To improve the success of transferring those engineered APs from model species to crops we must understand how they will interact with metabolism and how the plant physiology affects their performance. Here we used multiple mathematical modelling techniques to analyse and compare existing AP designs. We show that carbon-fixing APs are the most promising candidates to replace native photorespiration in major crop species. Our results demonstrate the different metabolic routes that APs employ to increase yield and which plant physiology can profit most from them. We anticipate our results to guide the design of new APs and to help improve existing ones.

**Code:** https://gitlab.com/gain4crops/2024-paper

## Introduction

Rubisco is the primary site of carbon fixation in plants. Alongside CO_2_, ribulose-1,5-bisphosphate carboxylase/oxygenase (rubisco) can also react with oxygen to produce one molecule of 3-phosphoglycerate (3PGA) and one molecule of 2-phosphoglycolate (2PG). Through a process called photorespiration, the 2-carbon product of the rubisco oxygenase reaction is converted back to an intermediate that can replenish the Calvin-Benson-Bassham cycle (CBB cycle). In this process, CO_2_ and ammonia are released in the mitochondria and reassimilated in the chloroplasts. Photorespiration consumes ATP and reducing power and causes losses of approximately 26 % of fixed CO_2_ and up to 36 % of the yield of certain crops [1, 2]. Thus, next to efforts in reducing the oxygenase activity of rubisco, the carbon and energy efficiency of photorespiration can be improved [2, 3, 4, 5]. This is highly desired to feed an increasing human population and to mitigate climate change.

Alternative pathways (APs) can increase the carbon and energy efficiency of photorespiration while still serving the primary function of photorespiration: detoxifying 2PG and replenishing the intermediates of the CBB cycle following the rubisco oxygenase reaction. There are four ways this can be achieved, ranging from fixing an additional CO_2_ to two CO_2_ per cycle being released (Figure 1). For fixing additional carbon, the two carbon 2PG can be converted into a 3-carbon compound such as 3PGA by addition of one carbon from CO_2_. A carbon-neutral photorespiratory pathway, with no net loss or fixation of CO_2_, is accomplished by combining 2PG with other compounds to generate a 4- or 5-carbon compound that can replenish the CBB cycle directly. In partial decarboxylation pathways, such as native photorespiration, a 2PG molecule is decarboxylated to form a 1-carbon compound that can be combined with another 2PG to make a 3-carbon compound that can enter the CBB cycle. Finally, 2PG can be completely decarboxylated, releasing two CO_2_ that can be refixed by the CBB cycle.

**Figure 1:**
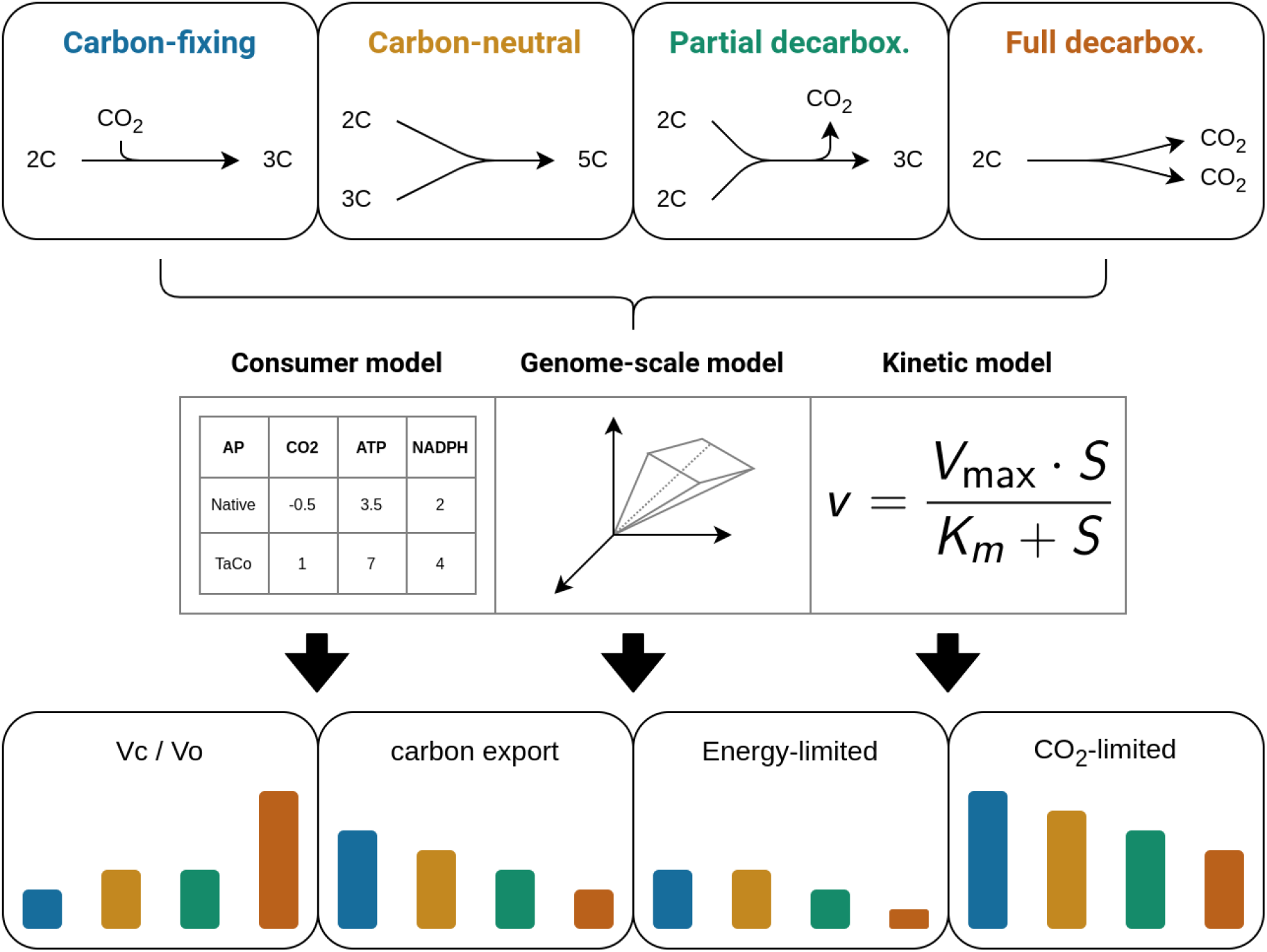
Graphical overview of this work. We analysed 12 alternative photorespiratory pathways (APs) that can be described using four general schemes regarding their carbon stoichiometry: carbon-fixing, carbon-neutral, partial decarboxylating and full decarboxylating pathways. The three modelling approaches we used are a simple stoichiometric consumer model, a genome-scale model solved using flux-balance analysis and a kinetic model to describe key reactions with higher fidelity. Modelling at these different scales allowed the mechanisms behind potential AP benefits to be evaluated.

From a carbon perspective an AP that fixes additional CO_2_ would be preferred. However, additional carbon fixation comes along with extra energy and redox requirements, decreasing the supply for other cellular processes. In contrast, decarboxylating APs generate reducing power that can be captured as redox equivalents as in the Kebeish and South AP3 pathways [6, 7]. In addition to that, decarboxylating APs can shift the rubisco activity towards carboxylation by releasing CO_2_ in the chloroplast and thereby increasing the local concentration of CO_2_. Thus, next to mere carbon efficiency, several other cellular parameters might determine the overall benefit of an AP.

Native photorespiration has cellular functions other than just clearing 2PG and replenishing the CBB cycle. Further functions include helping to balance ATP:NADPH supply and demand [8], increasing nitrogen assimilation into amino acids [9] and acting as an important source of 1C units [10] as well as serine and glycine [11]. An ideal APs must therefore still support these other roles of native photorespiration.

All these aspects together indicate that implementing photorespiratory APs to accomplish increased crop yield is far from trivial. In fact, simultaneously interfering with carbon, nitrogen, energy, and redox balances, means interfering with a highly complex, intertwined system. Thus, whether any change increases or decreases crop yield is dependent on metabolism, physiology and conditions, i.e. gas exchange differs with physiology and conditions and metabolism differs with conditions and growth stage/time of day etc. Towards rational engineering and full understanding of APs, global experimental analyses and mathematical modelling will likely be required to evaluate and understand the effect of variants. Exemplary for the challenge and for our yet limited understanding is the fact we still do not fully understand the true cause for the increased yields observed in complete decarboxylation pathways that have been experimentally tested in plants [12].

In this work, we quantitatively examine 12 APs using different mathematical approaches (Figure 1), building on previous modelling efforts [4]. These APs include those that have experimentally been tested *in planta*, but also include newly designed pathways that are either carbon-neutral or fix additional CO_2_ [13]. Our assessment includes carbon efficiency, energy and redox effects and effects emerging from altered subcellular CO_2_ levels in order to make judgments on the yield increases that one can expect on the system level. In addition, we make predictions on optimal variants of AP designs (e.g. optimal enzyme location) and which types of plants might benefit most from certain pathways and under which environmental conditions. By developing a mechanistic understanding of how APs can be effective, we identify the most promising pathways. Thereby, we hope to contribute to the challenge of increasing photosynthetic efficiency and thereby crop yield, to meet pressing societal issues.

## Results

### Description of APs

Several APs have been proposed or implemented to date, which can be classified with regards to their CO_2_ stoichiometry, ranging from fixing an additional CO_2_, via being CO_2_ neutral down to two molecules of CO_2_ being released (see Figure 2 and [4] for a review).

**Figure 2:**
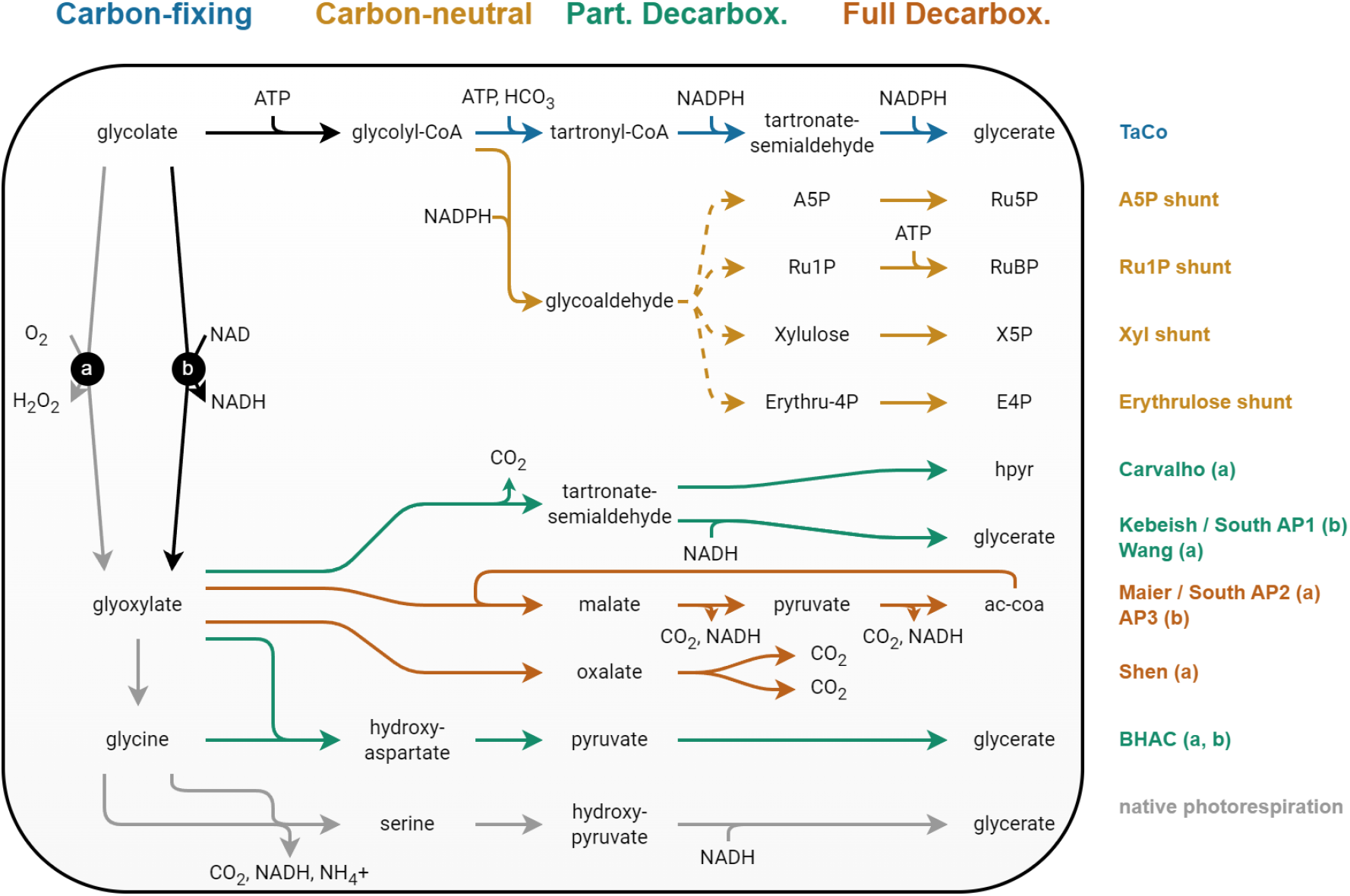
Schematic depiction of APs studied in this work. Grey arrows indicate wildtype reactions, colored arrows signify the classification regarding the carbon stoichiometry of the pathways: carbon-fixing, carbon-neutral, partial decarboxylating and full decarboxylating. Solid lines depict single reactions, while dashed lines depict intermediate steps that were omitted for clarity. The pathway variants **a** and **b** denote whether O_2_ / hydrogen peroxide (H_2_O_2_) or Nicotinamide adenine dinucleotide (NAD) / Nicotinamide adenine dinucleotide (reduced) (NADH) are used as the redox pair for glycolate dehydrogenase.

A CO_2_ fixing AP is the tartronyl-CoA pathway (TaCo), which activates glycolate with CoA, carboxylates the glycolyl-CoA to tartronyl-CoA and then reduces the tartronyl-CoA to generate glycerate (blue line in Figure 2) [13, 14]. The enzymes in the pathway and the metabolite tartronyl-CoA are not known to occur in nature, and therefore novel enzyme activities had to be engineered starting from related promiscuous enzymes. In particular, the glycol-CoA carboxylase, catalysing the fixation of bicarbonate to form tartronyl-CoA, has been subject to multiple rounds of engineering to alter the kinetic properties and to reduce the futile hydrolysis of ATP [14, 15]. While the TaCo pathway has been validated *in-vivo* [14], it has yet to be tested in photosynthetic organisms.

Carbon-neutral pathways have been proposed based on a glycolate reduction pathway, which converts glycolate to glycolaldehyde with glycolyl-CoA synthetase and glycolyl-CoA reductase, consuming ATP and NADPH (yellow lines in Figure 2) [13]. The glycolaldehyde is subsequently condensed with a sugar phosphate to generate longer chain sugars or sugar phosphates that can re-enter the CBB cycle via further conversion steps. Four variants have been proposed depending on the intermediates generated, namely arabinose-5P, ribulose-1P, erythrulose or xylulose. These carbon-neutral pathways required the engineering of novel enzyme activities for the two steps of the glycolate reduction. These novel pathways have been validated *in-vivo* [13], but are yet to be tested in photosynthetic organisms.

Partial decarboxylating pathways rely on combining two glycolate molecules and releasing one CO_2_ to generate a three-carbon intermediate (green lines in Figure 2). Several pathways with this design have been experimentally validated in plants. The pathways can be divided into those that rely on the *E. coli* glycolate catabolic pathway of glyoxylate condensation via tartronic semialdehyde (Carvalho, Kebeish/South AP1 and Wang pathways) [5, 6, 7, 16] and the β-hydroxyaspartate cycle (BHAC), which is the primary glycolate assimilation pathway in marine proteobacteria [17, 18]. The tartronic semialdehyde-based pathway generates glycerate that can be phosphorylated and re-enter the CBB cycle. The Kebeish/South AP1 and Wang pathways have been implemented in chloroplasts and the Carvalho pathway has been expressed in the peroxisome. The BHAC pathway generates a 4C product from two glycolate molecules, oxaloacetate (OAA), which can directly be incorporated into biomass via aspartate or can be used to replenish the CBB cycle. For this it must be decarboxylated to generate a three carbon CBB cycle intermediate, for instance 3PGA, via phosphoenolpyruvate-carboxykinase and enolase. So far the BHAC pathway has been implemented in peroxisomes of Arabidopsis [17]. The BHAC pathway could also theoretically be expressed in the plastid, which would facilitate CO_2_ reassimilation, but this remains to be tested in plants. The partial decarboxylation APs all release 0.5 CO_2_ per rubisco oxygenase reaction, the same as in native photorespiration, but are proposed to be more energy efficient as they all avoid the release and subsequent refixation of ammonium.

Complete decarboxylation pathways convert glycolate to CO_2_ using either malate synthase and pyruvate dehydrogenase (Maier/SouthAP2/AP3) [7, 19], or oxalate oxidase (Shen)[20] (dark orange lines in Figure 2). The primary difference between the two options is the fate of reductant released from glyoxylate oxidation. Malate synthase generates NADPH and pyruvate dehydrogenase generates NADH, whereas in the oxalate oxidase pathway two redox equivalents are transferred to water generating two H_2_O_2_. The CO_2_ released by glycolate decarboxylation can re-enter the CBB cycle and therefore complete decarboxylating pathways can still fulfil the requirement of converting 2PG into a CBB cycle intermediate.

In both partial and complete decarboxylating APs and in native photorespiration, oxidation of glycolate is the first step (Figure 2 a, b). Variants exist on in which subcellular compartment this oxidation takes place, and on what the final electron acceptor is of the oxidation. In native photorespiration, the Carvalho pathway and BHAC (perox.) pathway, glycolate oxidase is located in the peroxisome and converts glycolate to glyoxylate generating H_2_O_2_. In the Maier/South AP2, Shen, and Wang pathways [5, 7, 19, 20], glycolate oxidase was relocated to the chloroplast potentially altering subcellular redox dynamics. Glycolate dehydrogenase from E. coli, which generates NADH, was used for the Kebeish and South AP1 pathways and glycolate dehydrogenase from the algae Chlamydomonas reinhardtii (CrGDH), with an as yet unknown electron acceptor, was used in the South AP3 pathway [7, 21]. The different electron acceptors of the glycolate oxidation reaction have effects on the energy state and redox metabolism in the chloroplast, potentially affecting stress responses to high-light.

Overall, the above-described APs all convert 2PG to a CBB cycle intermediate. Yet, they differ in the CO_2_ stoichiometry, the energetic and redox costs, and their subcellular location.

### A consumer model allows comparison of APs stoichiometries

To compare the ATP, redox equivalents and CO_2_ stoichiometry of both the CBB cycle, photorespiration and APs it is convenient to define the pathways as closed cycles that regenerate one molecule of ribulose-1,5-bisphosphate (RuBP) and fix or release CO_2_ and triose phosphates in a so-called consumer model [13]. In this way, photorespiration can be separated from the CBB cycle to directly compare energy and CO_2_ stoichiometries. To calculate the ATP, redox equivalents and CO_2_ costs of the APs the individual reaction steps starting and ending at RuBP were summed (Table 1 and supplementary file 1).

**Table 1:** Consumer model allows comparison of APs stoichiometries. ATP, redox equivalent and CO_2_ stoichiometry of AP defined as converting 2PG to RuBP (extended from [4]). Positive values represent consumption, negative values production. The total energy cost assumes 2.5 ATP per redox equivalent. To account for differences in CO_2_ stoichiometry, the energy costs for 0 net change in CO_2_ were calculated by assuming the CBB cycle can compensate for the release / uptake of CO_2_ caused by photorespiration or APs. For example, for native photorespiration an additional 1.5 ATP and 1 redox equivalent must be spent to compensate for the 0.5 CO_2_ released. For the TaCo pathway 3 ATP and 2 redox equivalents are spared as 1 CO_2_ is already fixed by the AP.

| AP | CO<sub>2</sub> | ATP | Redox eq. | Total energy cost | ATP at net-zero CO<sub>2</sub> | Redox eq. at net-zero CO<sub>2</sub> | Total energy cost at net-zero CO<sub>2</sub> |
| --- | --- | --- | --- | --- | --- | --- | --- |
| TaCo | 1 | 7 | 4 | 17 | 4 | 2 | 9 |
| Ara5P/Ru1P shunts | 0 | 4 | 2 | 9 | 4 | 2 | 9 |
| Xyl./Eryth. shunts | 0 | 5 | 2 | 10 | 5 | 2 | 10 |
| BHAC (plastid) | -0.5 | 3 | 1 | 5.5 | 4.5 | 2 | 9.5 |
| Kebeish/South AP1 | -0.5 | 3 | 1 | 5.5 | 4.5 | 2 | 9.5 |
| BHAC (perox.) | -0.5 | 3 | 2 | 8 | 4.5 | 3 | 12 |
| Carvalho | -0.5 | 3 | 2 | 8 | 4.5 | 3 | 12 |
| Wang | -0.5 | 3 | 2 | 8 | 4.5 | 3 | 12 |
| Photorespiration | -0.5 | 3.5 | 2 | 8.5 | 5 | 3 | 12.5 |
| South AP3 | -2 | 2 | -2 | -3 | 8 | 2 | 13 |
| Maier/South AP2 | -2 | 2 | -1 | -0.5 | 8 | 3 | 15.5 |
| Shen | -2 | 2 | 1 | 4.5 | 8 | 5 | 20.5 |
| CBB cycle | 1 | 3 | 2 | 8 |  |  |  |

The three complete decarboxylation pathways, South AP3, Maier/South AP2 and Shen, require the same amount of ATP but differ in their redox equivalent stoichiometry, e.g. production of H_2_O_2_ or NAD(P)H (Table 1). Here, and in subsequent analyses, we assumed the Chlamydomonas reinhardtii glycolate dehydrogenase (CrGDH) used in the South AP3 pathway indirectly produces NADH, although the exact electron acceptor remains unknown [21]. The South AP3 and Maier/South AP2 pathways have the lowest total energy costs because they net generate NAD(P)H from the decarboxylation of glycolate (Table 1). Compared to native photorespiration, the partial decarboxylation pathways BHAC (perox.), Carvalho, and Wang pathways save 0.5 ATP and 0.5 NADPH by avoiding ammonia release, but require an additional NADH for reduction of tartronic-semialdehyde and do not gain 0.5 NADH from the oxidation of glycine. Therefore, avoiding the cost of ammonium release in these pathways only saves 0.5 ATP per rubisco oxygenase conversion when compared to native photorespiration. The Kebeish/South AP1 and BHAC (plast.) pathways have the same benefits from avoiding ammonia release but also produce an additional NADH from the use of glycolate dehydrogenase in place of the glycolate oxidase used by the Carvalho, Wang, and BHAC (perox) pathway. The carbon-neutral pathways require 0.5-1.5 more ATP than native photorespiration but require the same amount of redox equivalents (Table 1). The carbon-fixing TaCo pathway has the largest total energy cost as redox equivalents and ATP are required to fix CO_2_ in this pathway (Table 1). Overall, there is a positive correlation between CO_2_ released/taken up and the total energy cost of the pathway. Here, carbon-fixing pathways have the largest energy cost and CO_2_-releasing pathways, which can generate redox equivalents from the oxidation of previously fixed carbon, have the lowest energy cost.

### Net-zero CO_2_ consumer model accounts for the cost of CO_2_ release

The previous consumer model does not account for the increase in energetic cost caused by CO_2_ release, which needs to be refixed by the CBB cycle, or the reduction in energetic costs if CO_2_ is fixed by the APs. Accounting for the cost of modified CO_2_ stoichiometry provides a different perspective and allows for more direct comparison between pathways. For example, in native photorespiration if we also account for the cost of fixing the additional 0.5 carbons required to compensate for the loss of CO_2_ by glycine decarboxylase, then an additional cost of 1.5 ATP and 1 redox equivalent are required (with the simplifying assumption that the CBB cycle is able to fix CO_2_ with no associated rubisco oxygenase or photorespiration). Therefore, the total cost for native photorespiration assuming no net release/uptake of CO_2_, is 5 ATP and 3 redox equivalents per regeneration of RuBP (Table 1). Similar calculations were performed for all APs. Note that for the TaCo pathway, which fixes one CO_2_, we calculated a benefit in terms of ATP and NADPH spared by not requiring CBB cycle flux.

The complete decarboxylation pathways now have the largest total energy costs as an additional 6 ATP and 4 NADPH are required to refix the 2 CO_2_ that are lost by these pathways (Table 1). The partial decarboxylation pathways have a small additional energy cost for refixation of 0.5 CO_2_ and the carbon-neutral pathways have no additional costs/benefits as no additional CO_2_ must be fixed. In contrast, the TaCo pathway spares 3 ATP and 2 NADPH as it fixes an additional CO_2_, reducing the total energy cost. Overall, from an energetic perspective the importance of accounting for CO_2_ stoichiometry becomes apparent as large direct energy costs in CO_2_-fixing pathways can be compensated for by the energy saving from additional CO_2_ fixation.

#### Stoichiometric network modelling accounts for system-wide effects of APs

While the consumer model offers a useful description of the pathway stoichiometries, it does not capture the interplay of APs with the wider metabolic network, such as amino acid metabolism. Therefore, we extended our analysis by integrating the individual APs in a largescale stoichiometric model of core plant metabolism based on an Arabidopsis leaf, including photosynthetic electron transport and subcellular compartments [22]. We used these models to investigate whether the APs can replace the complete flux of native photorespiration. Therefore, photorespiration was blocked by constraining the flux through glycine decarboxylase to zero. To ensure flux through photorespiration or an AP was required, we mimicked ambient CO_2_ partial pressure by fixing the rubisco carboxylase:oxygenase ratio at 3:1 [23].

### Carbon-fixing APs are more energy efficient

We first assessed the photosynthetic energy efficiency of the APs, which we defined as the CO_2_ fixed per photon absorbed. For this, a sink reaction for glyceraldehyde-3-phosphate (GAP) was fixed to 1 μmol s^−1^ and the optimisation objective set to minimisation of photon influx. Thus, the most energy efficient flux distribution was identified for each AP.

The photosynthetic energy efficiency of the APs can have largely different values compared to the WT, ranging from an 27 % increase in TaCo down to a -54 % decrease in the Shen pathway (Figure 3). The carbon-fixing TaCo pathway shows the largest increase, followed by the carbon-neutral Ara5P and Xylulose APs (Figure 3). The partial decarboxylation APs range from a small increase to almost no change and the complete decarboxylation APs show a large decrease in photosynthetic energy efficiency (Figure 3). In terms of energetic efficiency, the net carbon exchanged by the AP has the greatest effect, with carbon-fixing APs being the most energy efficient and decarboxylating APs being the least energy efficient (Figure 3). The second most important factor determining the relative energy efficiency is the fate of redox equivalents or electrons from 2PG. Conversion of 2PG to CO_2_ can generate three pairs of electrons which can be used to either reduce NAD(P)^+^ to generate redox equivalents or water to generate H_2_O_2_. APs which generate more NAD(P)H (Figure 2, Table 1 and supplementary file 1) therefore have an advantage in terms of energy efficiency (Figure 3).

**Figure 3:**
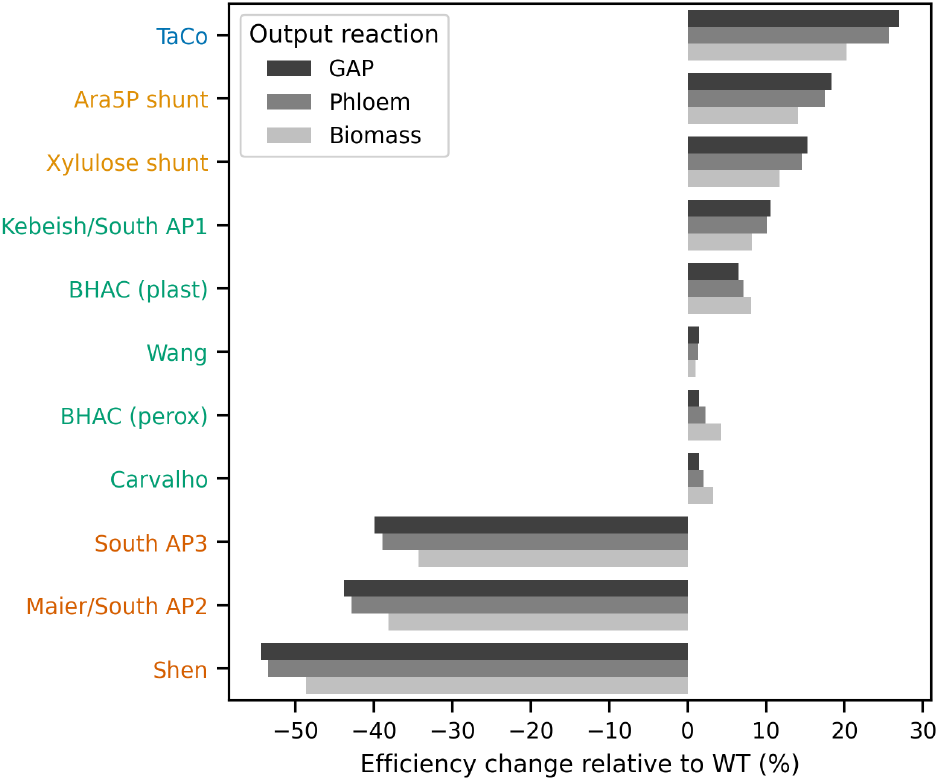
Change in photosynthetic energy efficiency (CO_2_ fixed per photon) of APs relative to WT photorespiration for a rubisco carboxylase:oxygenase ratio fixed at 3:1 calculated by stoichiometric FBA modelling. Negative numbers denote a decrease in CO_2_ fixed per photon relative to the WT, while positive numbers denote an increase. Model outputs were fixed to either glyceraldehyde 3-phosphate (GAP), phloem exudate or biomass represented by grey shading. AP label colours signify the amount of CO_2_ the AP fixed or released by the alternative pathways; blue, 1; yellow, 0; green, -0.5; orange, -2. The absolute CO_2_ fixed per photon range from 13.63 for GAP output to 17.67 for biomass, see supplementary Figure S2.

### APs can provide biosynthetic energy efficiency benefits

Photorespiration interacts with other metabolic pathways besides the CBB cycle. For example, carbon can be withdrawn from native photorespiration to provide one carbon units (CH_2_-THF), glycine or serine which can result in release of less than 0.5 carbons per rubisco oxygenase reaction [11, 24, 25, 26, 27, 28]. Similarly, intermediates can also be withdrawn from APs and used to synthesise amino acids or other biomass precursors.

To calculate the impact that APs have on wider metabolism, we calculated the relative energy efficiency in the presence of APs with more complex cellular outputs including Arabidopsis phloem exudate or biomass [29, 30]. In general, for the carbon-fixing, carbon-neutral, and Kebeish/South AP1 pathways the benefit of the APs decreases as the output complexity increases (Figure 3). Phloem exudate and biomass contain amino acids which can already be efficiently synthesised by native photorespiration. Therefore, the benefit of these APs is smaller when the cell is producing amino acids compared to GAP alone because these APs do not generate amino acids or amino acid precursors as intermediates. In contrast, the BHAC and Carvalho pathways demonstrate an enhanced benefit when synthesising biomass or phloem exudate compared to GAP alone (Figure 3). Synthesis of amino acids with five-carbon backbones is more efficient in the presence of the BHAC and Carvalho pathways as additional separate decarboxylation steps and subsequent refixation of CO_2_ are not required to synthesis these amino acids. Similarly, the complete decarboxylation pathways also show a benefit (i.e. decreased energy efficiency penalty) when synthesising biomass or phloem exudate in comparison to just GAP alone (Figure 3). Specifically, in the Maier/South AP2 and South AP3 pathways, carbon can be withdrawn as malate from malate synthase. Malate can then be used as the carbon backbone for generating aspartate and other derived amino acids. In this way, carbon is conserved and not released as CO_2_ that must be refixed with an associated energetic cost.

APs can have some small additional benefits compared to native photorespiration when cells are synthesising biomass, and these will depend on the precise amino acid demands of the leaf. The WT model predicts here that only 6 % of the carbon entering native photorespiration is withdrawn as serine, substantially less than the 32 % reported for Tobacco leaves [11]. All the APs investigated here bypass the serine producing steps of native photorespiration and could therefore disadvantage the plant if serine is in high demand and other serine producing pathways are unable to compensate. Overall however, the greatest effect of the alternative pathways is on the efficiency of carbon fixation and any changes in the energy efficiency of synthesising biomass precursors is likely to be relatively minor.

### APs alter the ATP and NADPH demand

The linear electron flow of photosynthesis supplies a fixed stoichiometry of ATP and redox equivalents that does not necessarily match the demand of the cell. Plants employ various mechanisms to match the supply to the demand, such as cyclic electron flow, but imbalances can potentially lead to photosynthetic inefficiency or damage [8]. Introducing APs can alter the ATP and redox demands of the cell with potentially positive or negative effects. We therefore quantified the ratio of ATP to NADPH demand of the cell in the presence of the APs to see whether the demand is shifted towards or away from the ratio supplied by linear electron flow. For this we identified the most energy-efficient flux distribution by forcing a GAP outflux of 1 μmolm^−2^s^−1^ while minimising the photon input and then quantified the fluxes through ATP synthase and ferredoxin-NADP reductase to quantify the net ATP and reductant demand. Because the model represents autotrophic leaf metabolism these reactions are the primary source of ATP and NADPH and can therefore be used to calculate the net ATP:NADPH demand of the cell.

Here, only the Carvalho, Wang and peroxisomal BHAC pathways, with an ATP:NADPH demand of 1.5, are closer to the supply from linear electron flow (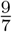 or 1.28) than that of a plant with WT photorespiration (1.56) (Figure 4). The lower ATP:NADPH demand in the Carvalho, Wang and BHAC (perox.) pathways is due to avoiding the need for ammonia reassimilation and the associated ATP cost. APs that capture the redox equivalents from complete glycolate decarboxylation as NADPH, such as the Maier and South AP3 pathways, significantly increase the demand of ATP relative to NADPH (up to 2.75 in South AP3) (Figure 4). This is caused by the increased NADPH supply from glycolate decarboxylation which decreases the NADPH that must be supplied from photosynthetic electron flow. In contrast, the Shen pathway, which also completely decarboxylates glycolate, does not capture the redox equivalents as NADPH and instead produces H_2_O_2_, resulting in an ATP:NADPH demand of 1.57 (Figure 4).

**Figure 4:**
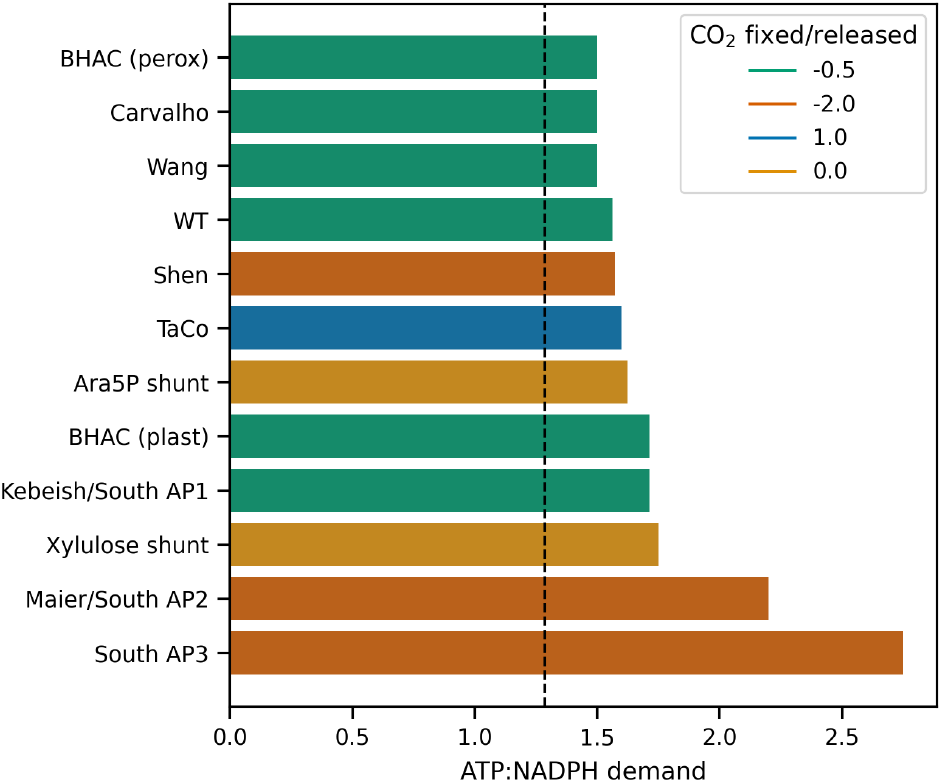
APs alter the ATP:NADPH demand of carbon fixation. The net ATP:NADPH demand of carbon fixation to glyceraldehyde 3-phosphate in the presence of APs calculated using the stoichiometric model and FBA, and assuming a rubisco carboxylase:oxygenase activity of 3:1. Dashed line represents the ATP:NADPH supplied by linear electron flow through the photosystems 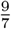.

Overall, most of the APs cause relatively minor changes in the ATP:NADPH demand compared to native photorespiration with the notable exception of the complete decarboxylating Maier/South AP2 and South AP3 pathways which cause large increases in the ATP:NADPH demand (Figure 4).

### Complete decarboxylation pathways require an increased rubisco carboxylation rate to have a benefit

Analysis so far has focused purely on photosynthetic energy efficiency and assumed no limitation of CO_2_ diffusion either from outside the cell or between subcellular compartments. However, CO_2_ diffusion poses a major limitation to photosynthesis in C3 plants [31]. The complete decarboxylation pathways decrease the photosynthetic energy efficiency relative to WT photorespiration if unlimited CO_2_ diffusion is assumed. However, if CO_2_ diffusion into the chloroplast is limited, releasing CO_2_ specifically within the chloroplast and thereby increasing the carboxylation rate relative to the oxygenation rate could be beneficial. We therefore determined the increase in carboxylation rate relative to the oxygenation rate necessary to compensate for the increased energy cost of the complete decarboxylation pathways.

To model the effect of an increased chloroplast CO_2_ concentration, we constrained the rubisco carboxylase:oxygenase ratio of rubisco to a range of values and calculated the photosynthetic energy efficiency relative to the WT in terms of CO_2_ fixed per photon absorbed.

At a rubisco (rubisco carboxylase:oxygenase) ratio of 3:1 the complete decarboxylation pathways were less energetically efficient than WT photorespiration (Figure 5). However, these pathways can become more energetically efficient than WT photorespiration if they are able to increase the chloroplast CO_2_ concentration such that the rubisco (rubisco carboxylase:oxygenase) ratio reaches 4.8-6.1:1 (Figure 5). This represents a 60-103 % increase relative to the rubisco carboxylase:oxygenase ratio of 3:1 assumed for native photorespiration. The remaining APs that release 0.5 CO_2_, are carbon-neutral or fix additional CO_2_, continue to show an increased photosynthetic efficiency even at lower rubisco carboxylase:oxygenase ratios (Figure 5).

**Figure 5:**
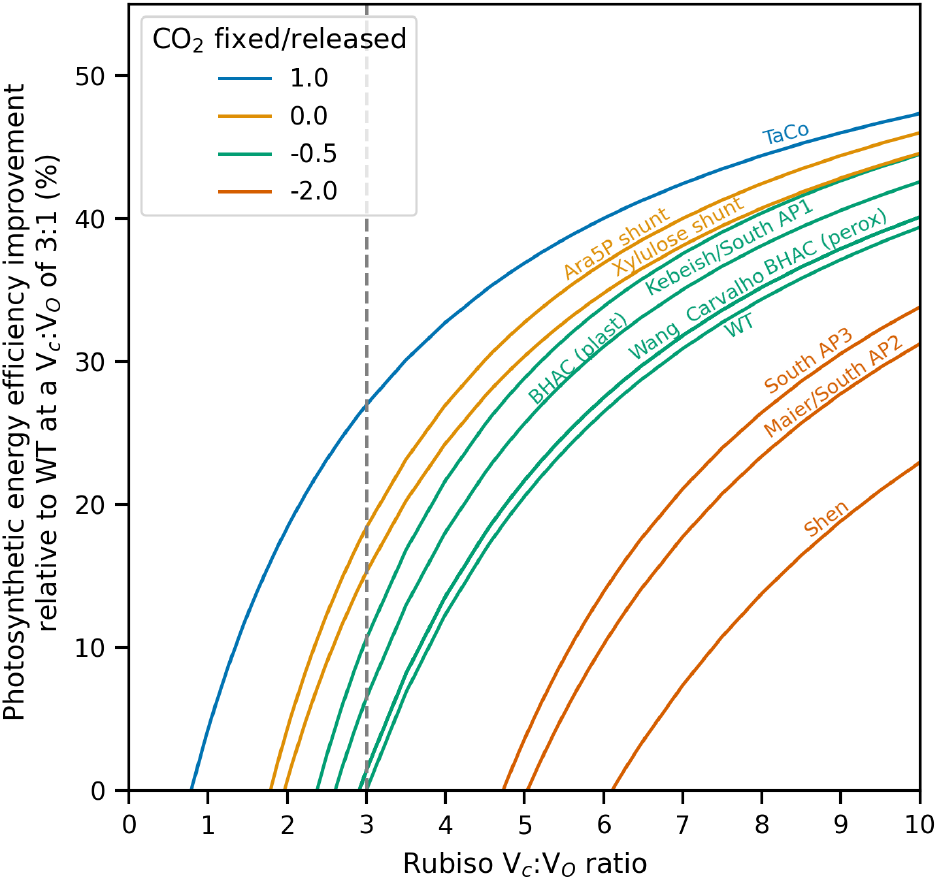
Complete decarboxylation pathways require an increased rubisco carboxylase:oxygenase ratio to achieve an increased photosynthetic energy efficiency relative to WT photorespiration. Effect of altered rubisco carboxylase:oxygenase ratio on photosynthetic energy efficiency of APs relative to WT photorespiration at a fixed rubisco carboxylase:oxygenase ratio of 3:1 (dashed line). Simulations were performed using the stoichometric model and FBA with the objective set to maximisation of glyceraldehyde 3-phosphate production. Photosynthetic energy efficiency was defined as the CO_2_ fixed per photon absorbed.

### Native CO_2_ refixation capacity affects AP benefit

Another factor that needs to be taken into account is that in native photorespiration CO_2_ set free by glycine decarboxylase activity in the mitochondria can diffuse out of the cell before it reaches the chloroplast, and is therefore only partly refixed. Plants have evolved mechanisms to recapture this CO_2_ that would otherwise be lost, such as placing chloroplasts around the cell periphery [28, 32, 33, 34, 35] or relocating the decarboxylation step to the bundle sheath cells, as in C3-C4 intermediate photosynthesis [36, 37]. The exact amount of CO_2_ refixation depends on both the plant species and the environmental conditions [32, 37]. APs can relocate the site of CO_2_ release from the mitochondrion to the chloroplast and therefore potentially increase the proportion of refixed CO_2_.

CO_2_ diffusion is dependent on a series of resistances between the external and internal airspace and between subcellular organelles [28, 33, 34, 35]. These resistances were simplified in our stoichiometric model to a constraint on the fraction of CO_2_ released in mitochondria that can be refixed by chloroplasts (refixation potential), with the remainder assumed to exit the leaf. We modelled the effect of different CO_2_ refixation potentials in the WT plant on the relative carbon export of plants expressing APs compared to the WT. We assumed that relocation of CO_2_ release by the APs to the chloroplast results in complete recapture of CO_2_ released by the APs, therefore representing the maximum potential benefit. The input of CO_2_ from outside the cell was fixed to the WT value to represent a CO_2_ diffusion limited condition and prevent cells from compensating for CO_2_ lost from photorespiration or APs by simply importing more CO_2_. Photon input was constrained to the WT value representing an energy limited condition and the optimisation objective set to maximisation of GAP production.

In general, as the CO_2_ refixation potential of the WT plant increases, the benefit of an AP which recaptures this otherwise lost CO_2_ decreases (Figure 6A). The maximum carbon export increase of any AP that recaptures otherwise lost CO_2_ is 20 % for a rubisco carboxylase:oxygenase ratio of 3:1 (Figure 6A). Under the energy limited condition modelled here, the APs that are more energetically efficient than native photorespiration (carbon-fixing, carbon-neutral and partial decarboxylation pathways) can all achieve this maximum 20 % benefit (Figure 6A “All other APs”). In contrast, the complete decarboxylation pathways, which are less energy efficient than native photorespiration, fixed less CO_2_ than the WT plant at all CO_2_ refixation potentials (Figure 6A, dashed orange lines). Next we evaluated whether residual flux through the native photorespiratory pathway could be beneficial in combination with an AP. Native photorespiration potentially loses CO_2_ from the cell by releasing it in the mitochondria, affecting the carbon export. APs can recapture this CO_2_ by releasing it in the chloroplast, but can be more energetically expensive. Therefore, for the complete decarboxylation APs, there is a tradeoff between the relatively more energy efficient native photorespiration and the more carbon efficient AP. We therefore repeated the previous analysis but allowed unlimited flux through native photorespiration and identified the optimal flux through native photorespiration and the AP under a CO_2_ and energy limitation.

**Figure 6:**
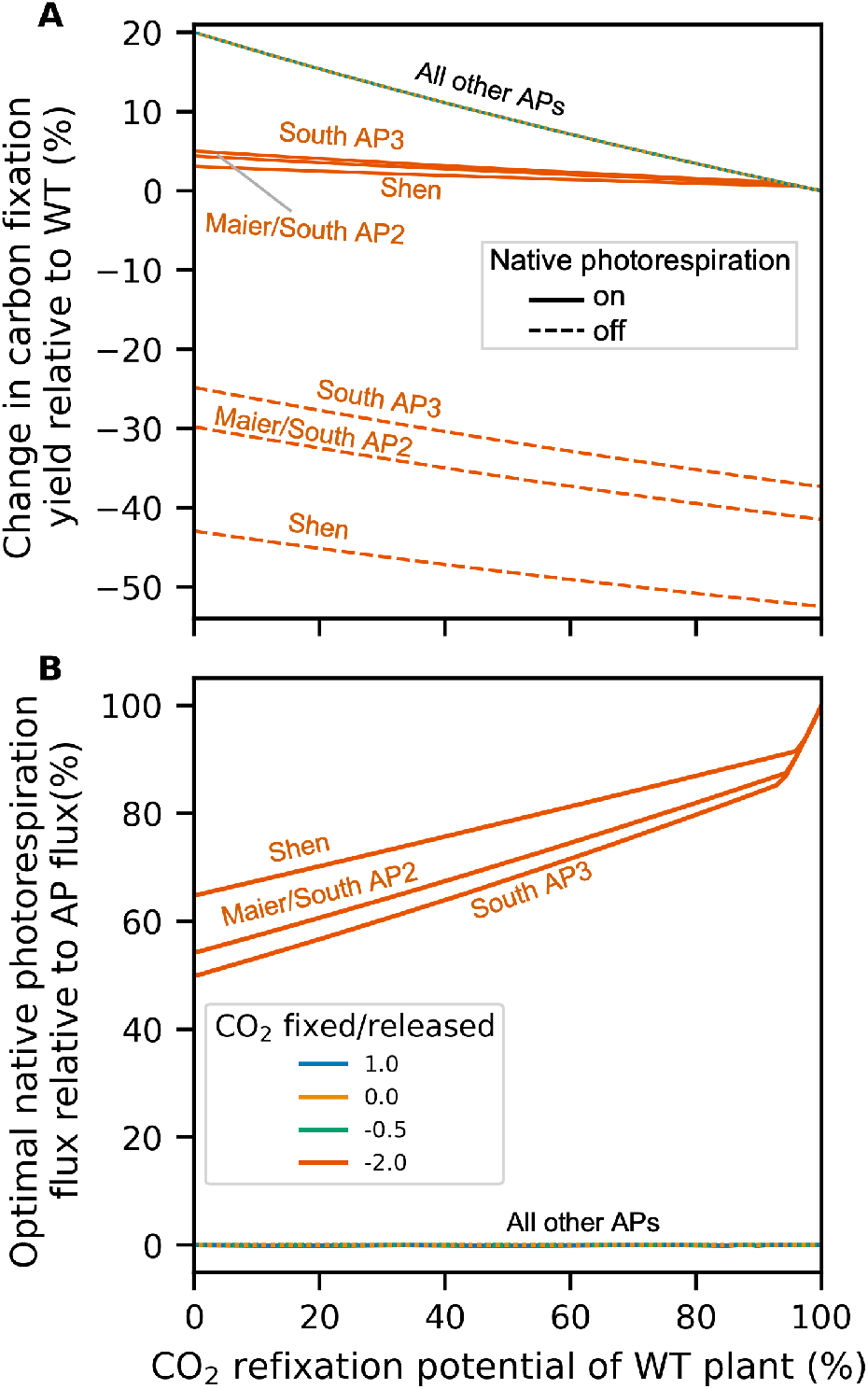
The benefit of an AP that recaptures otherwise lost CO_2_, depends on how effective the WT plant is at already refixing CO_2_ released by photorespiration. A) The effect of CO_2_ refixation in WT plants on the relative benefit of photorespiratory APs to carbon fixation calculated using the stoichiometric model and FBA. Refixation potential is defined as the fraction of CO_2_ released from photorespiration in the mitochondria that could enter the chloroplast rather than leaving the cell. The refixation potential of a WT cell was fixed between 0-100 % and the objective set to maximisation of glyceraldehyde 3-phosphate production (GAP). Rubisco carboxylase:oxygenase activity was constrained to 3:1. When simulating the APs, photon and CO_2_ inputs were constrained to the WT value required to generate GAP at a rate of 1 μmol s^−1^ representing an energy and CO_2_ limited condition. Dashed lines are with flux though native photorespiration blocked; solid lines are with native photorespiration free to carry flux. B) The optimal flux through native photorespiration relative to the AP at varying CO_2_ refixation potentials. Photon and CO_2_ inputs were constrained as in A. 100 % means that there is no flux through the AP and 0 % means that the AP completely replaces photorespiration.

Under these energy- and CO_2_-limited conditions, the optimal flux distribution for the more energy efficient pathways required zero flux through photorespiration (Figure 6B). In contrast, for the less energy efficient complete decarboxylation pathways, the optimal amount of native photorespiratory flux varied with the CO_2_ refixation potential (Figure 6 B). In other words, residual flux though native photorespiration can compensate for the energy inefficiency of complete decarboxylation pathways while the APs act to recapture otherwise lost CO_2_. Therefore, some residual photorespiratory flux can be beneficial in the presence of complete decarboxylation pathways, particularly when the CO_2_ refixation potential is low (Figure 6 A, solid orange lines). The optimal flux through native photorespiration in combination with an AP may vary dynamically with the environmental conditions which can affect CO_2_ diffusion and the refixation potential of the plant. Overall, assuming approximately 25 % of CO_2_ is refixed in a WT plant, all AP designs are advantageous when both CO_2_ and energy are limiting, and when operating in combination with native photorespiration (Figure 6A, solid lines).

### Understanding CO_2_ dynamics requires kinetic modelling

Stoichiometric modelling demonstrated the energetic and stoichiometric benefits of the AP designs, how flux through certain APs can support amino acid biosynthesis and the potential beneficial effects of avoiding CO_2_ release in the mitochondria. However, the stoichiometric model required fixing the rubisco carboxylase:oxygenase ratio, whereas *in-vivo* it depends on the concentrations of O_2_ and CO_2_ which can vary dynamically. Therefore, to model the effect of varying CO_2_ diffusion and CO_2_ concentration more accurately, we developed a kinetic model of photosynthesis to test the different AP designs.

The kinetic model was developed as a system of ordinary differential equations by combining models of the CBB cycle [38] and photorespiration [39], as well as a complete description of rubisco kinetics including carboxylation and oxygenation reactions [40]. CO_2_ was modelled as a dynamic variable to capture the effect of the various APs on the CO_2_ concentration in the chloroplast. A fixed proportion (25 %) of CO_2_ released by glycine decarboxylase in the mitochondria was assumed to diffuse back into the chloroplast [32]. The APs were implemented on top of this WT model and are grouped by their CO_2_ stoichiometry, with the best performing variant of each group shown in the following results. For a complete description of the model as well as a comparative analysis of intermediate model stages see supplementary material section S2.

### Carbon-fixing pathways are more efficient at exporting carbons

We first evaluated the different AP designs at reference conditions of 400 ppm atmospheric CO_2_ and an illumination of 700 μmol/m^2^s. From this analysis we calculated the rubisco carboxylation and oxygenation rates (V_*c*_ and V_*o*_) and rubisco carboxylase:oxygenase ratio. We further extended the evaluation by considering export of triose-phosphates and hexoses as a proxy for plant growth and defined the carbon-use efficiency as the ratio of carbon export rate relative to the rubisco carboxylase rate.

The carbon-fixing pathways show a reduced rubisco carboxylation rate (less than 75 % of the WT) and slightly increased oxygenation rate, leading to a reduced rubisco carboxylase:oxygenase ratio relative to the WT (Figure 7). The complete decarboxylation APs show the exact opposite, with a higher carboxylation rate (more than 125 % of the WT) and lower oxygenation rate, leading to a rubisco carboxylase:oxygenase ratio of more than 200 % relative to the WT (Figure 7). Even if the secondary carboxylation steps like glycolyl-CoA carboxylase (GCC) in the carbon fixing TaCo pathway are counted in the total carboxylation rate, this general trend stays the same, even though now the total carbon fixation of the carbon-fixing APs is slightly increased (Figure 7, transparent area). Contrastingly, the carbon export is highest for the carbon-fixing APs (around 120 %) and lowest for the complete decarboxylation APs at slightly above 100 % (Figure 7). This is reflected by the carbon-use efficiency, which is highest for the carbon-fixing APs at around 190 % and lowest for the complete decarboxylation APs at around 80 % (Figure 7). This increase is carbon-use efficiency in carbon-neutral and carbon-fixing APs is caused by the fact that they generate intermediates, other than CO_2_, that can enter the CBB cycle. By increasing the input of intermediates into the cycle, more carbon can be withdrawn without requiring additional fixation of CO_2_ by rubisco. Therefore, carbon-fixing and carbon-neutral alternative photorespiratory pathways increase the carbon use efficiency of the CBB cycle resulting in more carbon exported per rubisco carboxylase reaction.

**Figure 7:**
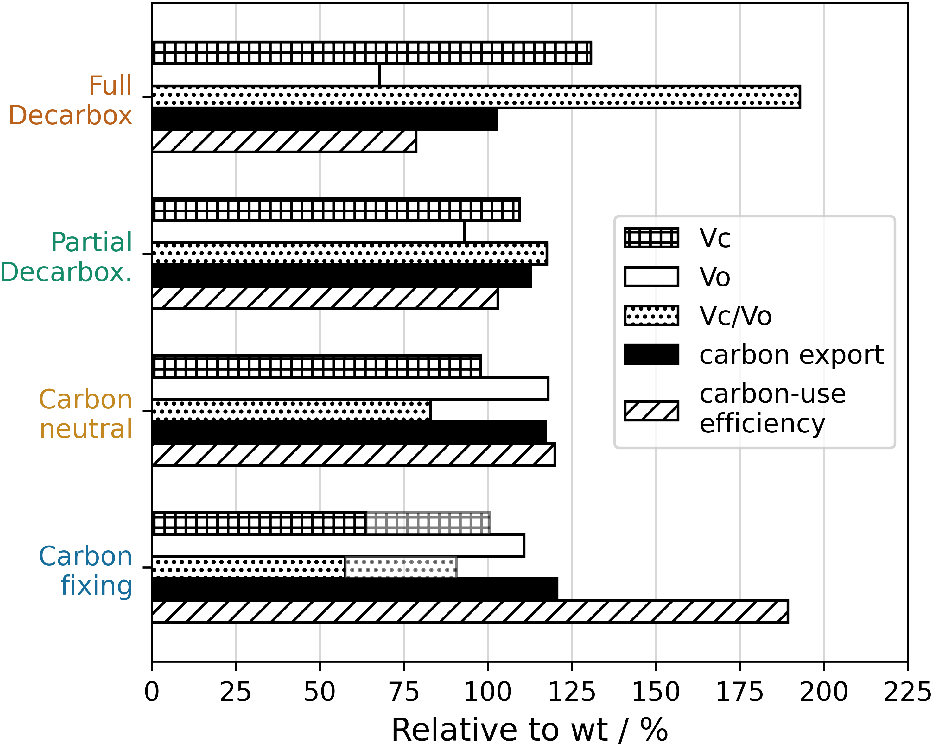
Steady-state performance indicators of plant metabolism simulated by the kinetic model for different AP designs, grouped by their CO_2_ stoichiometry. The simulation was performed at reference conditions of 400 ppm atmospheric CO_2_ and an illumination of 700 μmol*/*m^2^s. Transparent bars signify the addition of secondary carboxylation steps to the measurement of rubisco carboxylation rate for the carbon-fixing pathways. The different AP designs are grouped by their CO_2_ stoichiometry and only the pathway variant with the maximal carbon export of the group is shown.

### APs are most beneficial at low intracellular CO_2_

The activity and effectiveness of the APs, which we measure as the improvement of carbon export, depends on the local CO_2_ concentration in the chloroplast. The velocity at which this local CO_2_ can be replenished is dependent on the external CO_2_ concentration as well as the internal transport rate. We systematically evaluated all AP designs for both of these factors. The improvement of carbon export relative to the WT is highest for both low CO_2_-transport rate (Figure 8D), as well as low external CO_2_ concentration (Figure S1), in which the carbon-fixing pathways perform best (Figure 8D). Importantly, these are also the conditions where the absolute rates of carbon export are reduced (Figure 8B). The carbon-fixing APs consistently show the highest improvement relative to the WT, closely followed by the carbon-neutral APs and then by the partial decarboxylation APs (Figure 8D). The full decarboxylation APs show the lowest improvement and perform worse than the WT if the CO_2_-transport rate is increased more than 10 % (Figure 8 D). The improvement to carbon export diminishes for all APs under very high CO_2_ conditions (Figure S1).

**Figure 8:**
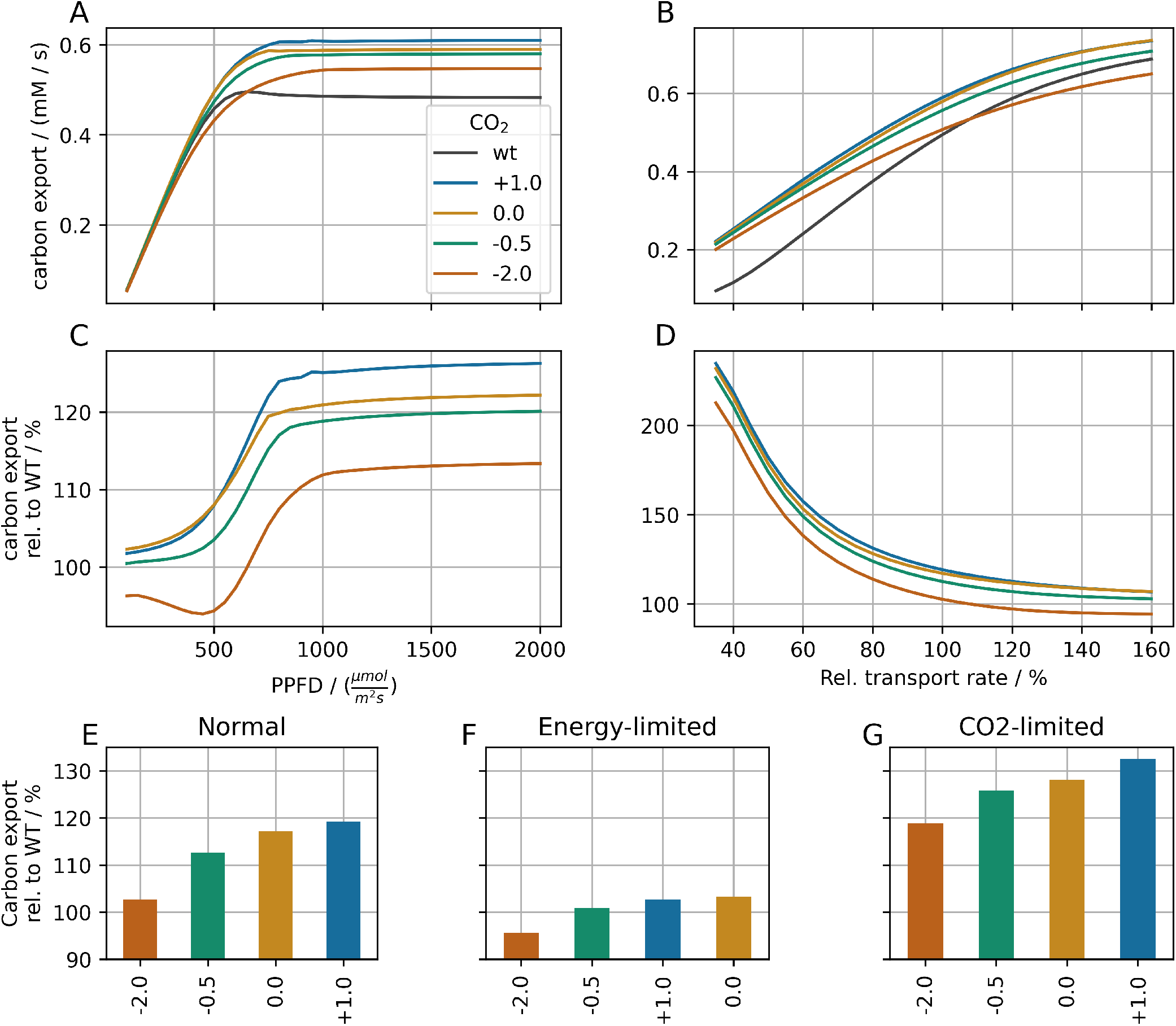
Steady-state carbon export of photorespiratory APs simulated by the kinetic model depending on photosynthetic photon flux density (PPFD) (A, C) or CO_2_ transport rate (B, D). Carbon export is shown in both absolute terms (A, B) and relative to the WT under the same conditions (C, D). Three representative scenarios are highlighted for a well-watered plant in a temperate climate with sufficient light (700 μmol*/*m^2^s) (E); a well-watered plant in temperate climate with low light (250 μmol*/*m^2^s), which is thus energy-limited (F); and a hot and dry climate with low water supply and thus partially closed stomata (90 % of WT CO_2_ transport rate), which is thus CO_2_-limited (G). The different AP designs are grouped by their CO_2_ stoichiometry (−2, -0.5, 0, 1) and only the pathway variant with the maximal carbon export of the group is shown.

### High-light increases AP benefit

All APs require more energy equivalents than the CBB cycle per carbon export and the full decarboxylation pathways even more than native photorespiration. Therefore, it is important to assess how much energy supply is necessary, such that the benefits of the pathways outweigh this energy cost. As light is the primary source of energy for plants, and natural light conditions are continuously changing, the AP performance should be assessed at a variety of light-conditions. We did this by systematically scanning the effect of illumination on the carbon export, by simulating the steady-state for each photosynthetic photon flux density (PPFD). Here, we found all APs show the highest relative increase in carbon export in high-light conditions, with the carbon-fixing APs showing the highest increase in carbon export at 125 % relative to the WT (Figure 8C). The second-best option are carbon-neutral pathways, which in low-light conditions outperform the carbon-fixing pathways (Figure 8 C illumination below 400 μmol). The complete decarboxylation pathways show a beneficial effect for medium and high-light conditions compared to the WT, but are disadvantageous in low light conditions compared to the other APs (Figure 8C). These results demonstrate that in medium to high-light conditions the benefits of all APs outweigh the increased direct energy cost, while at low light the increased energy demand outweighs the benefits for the complete decarboxylation APs.

### Environmental conditions require different AP designs

The previous results highlight that depending on the environmental conditions, which will affect CO_2_ diffusion and light intensity, different APs should be utilised to maximise carbon export. We predicted the carbon export improvement of different APs for three distinct scenarios. First, a normal scenario, which corresponds to a well-watered, plant in a temperate climate with sufficient light. Next, a scenario with a well-watered plant in temperate climate but low light, which is thus energy-limited. For this we used an illumination of 250 μmol/m^2^s. Lastly, a scenario of a hot and dry climate and high light, with partially closed stomata, which is thus CO_2_-limited. This we represented by lowering the carbon transport rate to 90 % of the WT.

In both the normal and CO_2_-limited scenario the carbon-fixing APs show the highest carbon export (Figure 8E, G). Under energy-limiting conditions carbon-fixing and carbon-neutral APs show similar carbon export (Figure 8F). In all cases, complete decarboxylation pathways show the lowest carbon export, with a negative effect in energy-limited conditions, a slightly positive effect in normal conditions and a positive effect in CO_2_-limited conditions (Figure 8E-G). Thus, in most scenarios the carbon-fixing APs are the preferred choice, while in energy-limited conditions the carbon-neutral APs might be more beneficial.

## Discussion

Using various mathematical models of plant metabolism, we have comprehensively identified and assessed the mechanisms by which APs can enhance carbon assimilation and growth. To understand how environmental conditions and plant physiology can affect pathway benefits, we also quantitatively assessed the effects of light intensity and CO_2_ availability on the different APs. We show that carbon-fixing APs, such as the TaCo pathway, have the greatest potential benefit over a range of conditions, and could provide an increase in carbon export from photosynthesis of over 20 %. Due to the initial exponential growth phase of plants, such a percentage could generate substantial gains in biomass over time [41].

### APs use distinct mechanisms to improve growth

Our work showed that alternative photorespiratory pathways can provide benefits to plants via five different mechanisms (Table 2): (i) An energy efficiency benefit by metabolising the 2PG produced by the rubisco oxygenase reaction in a way that uses less ATP and reducing power compared to native photorespiration. (ii) A biosynthetic energy efficiency benefit if intermediates of the alternative pathway can be used for biosynthetic reactions. (iii) Recapture of CO_2_ that could otherwise diffuse from the cell. (iv) An increased CO_2_ concentration at the site of rubisco in the chloroplasts caused by altered CO_2_ diffusion within the cell. (v) Increased carbon use efficiency of the CBB cycle caused by an additional input of CBB cycle intermediates, which enables more carbon to be exported per rubisco carboxylase reaction. As CO_2_ diffusion differs depending on cell morphology and leaf physiology, these CO_2_ related benefits can be different in different plant species. Thus, next to simply clearing the 2PG produced by the rubisco oxygenase reaction, alternative photorespiratory pathways can have broad effects on the plant in terms of energetics, biosynthesis and CO_2_ dynamics and these must be considered at the system level to evaluate the effectiveness of any pathway.

**Table 2:**
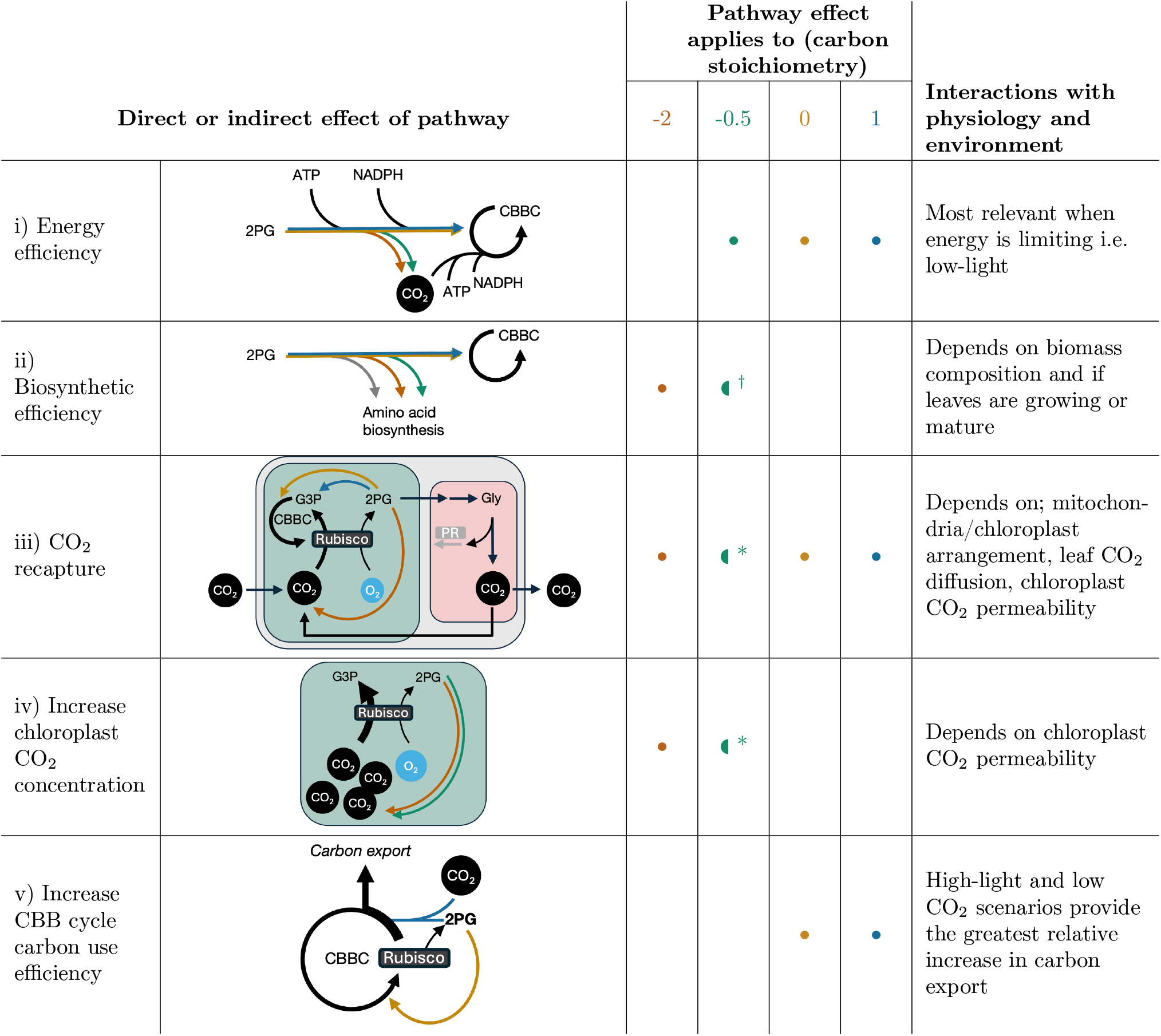
Alternative photorespiratory pathways can affect carbon assimilation and growth in multiple ways. *Only applies to pathways that relocate CO_2_ release to chloroplasts i.e not Carvalho or BHAC(perox.) pathways which are peroxisomal. ^†^Only applies to BHAC and Carvalho pathways not Wang, Kebeish/South AP1. 2PG, 2-phoshpoglycolate; CBBC, Calvin-Benson-Bassham cycle; G3P, glyceraldehyde 3-phosphate.

### Benefits of APs depend on environmental conditions

Importantly, the environmental conditions a plant grows in, such as light intensity, temperature, and water availability can affect the benefit of the different mechanisms that alternative pathways employ. Under low-light conditions, when energy is limiting, the energy efficiency of alternative pathways becomes critical to their effectiveness [12, 42]. Our models showed that alternative pathways that are more energy efficient than the native photorespiration will offer greater benefits in low-light, which is experimentally supported by growth enhancement in Arabidopsis expressing the Kebeish pathway and grown under low-light and short-day conditions [6]. In contrast, complete decarboxylation pathways, which are less energy efficient than native photorespiration, are predicted to show no benefits under these low-light conditions. Under high-light conditions energy efficiency no longer offers an advantage, and the pathways’ effects on CO_2_ diffusion and fixation become more important. As our analyses showed, all pathways perform best under high-light conditions (Figure 8A,C) suggesting that benefits from altered CO_2_ diffusion have the biggest potential to increase plant growth. This finding is also supported experimentally with complete decarboxylating pathways including South AP3, Shen, and Wang all showing increased benefits under high-light conditions [5, 7, 20]. Overall, the alternative pathways provide the greatest benefit over native photorespiration under high-light, and CO_2_-limiting conditions, with certain pathways also able to provide benefits under low-light conditions. The benefits of alternative pathways are comparable to those predicted for C3-C4 intermediate or C4 metabolism [43]. Indeed, the conditions which favour these naturally evolved mechanisms are also those we identified as beneficial for the alternative pathways. However, from a metabolic engineering perspective, the alternative pathways described here offer advantages over introducing C3-C4 or C4 metabolism into a C3 plant, as they can require as few as three genes compared to the >15 genetic modifications potentially needed to support the biochemistry, leaf anatomy, and intercellular transport of C3-C4 or C4 photosynthesis. Therefore, alternative pathways offer a more easily implementable solution to the same problems that C3-C4 intermediate and C4 photosynthesis have evolved to address.

### Applying new insights to previous results

By applying the insight we have gained, we can now try to better explain previous experimental observations. For example, complete decarboxylating pathways such as the Maier/South AP2, South AP3 and Shen by-passes experimentally showed growth benefits, but the molecular basis for these benefits was less clear, as previous computational analysis based on a kinetic model describing carbon fixation and subcellular CO_2_ conductance did not predict an enhanced rate of photosynthesis [12]. Here, we quantitively demonstrate that complete decarboxylating pathways can indeed enhance photosynthesis by increasing the CO_2_ concentration in the chloroplast and subsequently the rubisco carboxylase:oxygenase ratio, as well as by recapturing otherwise lost CO_2_. These effects are consistent with experimental measurements of unchanged or decreased CO_2_ compensation points in engineered plants expressing these pathways [7, 19, 20, 44]. Furthermore, for complete decarboxylating and some partial decarboxylating pathways, we demonstrated additional increases in energy efficiency in cells synthesizing amino acids for biomass synthesis or phloem exudate that have not previously been identified. Thus, through our work, we can now better explain the reasons why in previous studies a particular alternative pathway has generated benefits.

### Some aspects of APs require future investigation

Yet, despite improving our understanding of alternative photorespiratory pathways, a number of experimental observations remain unexplained. For example, the South AP3 pathway expressed in tobacco shows increased CO_2_ assimilation even at very high intracellular CO_2_ [7]. Decarboxylation pathways reduce rubisco oxygenase activity by increasing the local CO_2_ concentration at increased energetic costs. At very high intracellular CO_2_ this mechanism cannot offer any additional benefit, as CO_2_ concentration in the chloroplast should already be high. The mechanism of the benefit of South AP3 pathway even at high CO_2_ remains unexplained and may relate to the reported beneficial effects of expressing glycolate dehydrogenase alone, which could affect the efficiency of photosynthetic electron transport [6, 45, 46, 47]. Additionally, our models predicted that partial decarboxylating pathways should outperform full decarboxylating pathways across conditions. However, comparison of the partial decarboxylating South AP1 and the complete decarboxylating South AP3 expressed in tobacco in greenhouse trials, showed greater yield increases in the AP3 pathway [7]. In this context, it is important to realise that our model predictions reflect best-case scenarios, requiring optimal expression and kinetics of the pathway enzymes. While a discrepancy between our model predictions and experimental implementations of alternative pathways could point to shortcomings in our models, an alternative explanation for the discrepancy could be that an engineered plant might not yet have optimal enzyme expression levels. Thus, such discrepancy might also indicate potential for further improvements of plant performance.

## Conclusion

With this work, we have improved our mechanistic understanding of how alternative photorespiratory pathways can enhance carbon assimilation and growth, and used this to predict that the carbon fixing TaCo pathway represents the best option for increasing crop yields over a range of conditions. In future, crops, or specific cultivars, should be screened to identify those with limitations in CO_2_ refixation capacity or CO_2_ diffusion that make them most likely to benefit from engineering with alternative photorespiratory pathways. The models presented here could also be used to evaluate or develop new alternative pathway designs that may further increase yield gains e.g. by targeting specific growth scenarios, such as the juvenile or adult growth stages, or specific crop species. With the ability to more rationally engineer alternative photorespiratory pathways into suitable crops, and identify their optimal growing conditions, our work will hopefully contribute to realising the maximum impact of alternative photorespiratory pathways for improving crop yields.

## Methods

### Stoichiometric model

A stoichiometric model of core plant metabolism based on an Arabidopsis leaf, PlantCoreMetabolism_v3, was curated starting from a previously described model (PlantCoreMetabolism_v1_2, [22]). The model is available in SBML format as an XML file at https://gitlab.com/gain4crops/2024-paper. Python 3, COBRApy [48] and the CPLEX solver were used for FBA optimisations. All code required to reproduce the results is available at https://gitlab.com/gain4crops/2024-paper as Jupyter notebook files. Output fluxes to phloem exudate were defined from Arabidopsis phloem composition [29]. Output fluxes to biomass were defied as described in the AraGEM model [30]. Net ATP demands were calculated by quantifying the flux through both plastidic and mitochondrial ATP synthase. As the model is autotrophic, all ATP must ultimately be generated by either plastidic ATP synthase, or by mitochondrial ATP synthase using NADH generated in the plastid. To calculate the reductant demand the flux through proton pumping mitochondrial

NADH-dehydrogenase (which is used for generating ATP) was subtracted from the flux through plastidic ferredoxin-NADP reductase.

### Kinetic model

The ordinary differential equation (ODE) model was built mainly using two previously published models of the CBB cycle [38, 49] and photorespiration (PR) [39] and developed using Python-based software modelbase [50]. Rubisco kinetics including carboxylation and oxygenation reactions were described based on the rate equation from Witzel 2010 [40]. Energy metabolites, ATP and NADPH, were implemented as dynamic variables as in Matuszyńska 2019 [51] and their production modelled using a simplified light-dependent reaction with an additional quenching reaction to account for the different ATP and NADPH demands of the system. Thioredoxin based redox regulation of CBB cycle enzymes was linked to the energy status of the model via an NADPH-thioredoxin reductase reaction based on the description by Saadat 2021 [52]. CO_2_ was modelled dynamically with CO_2_ input from the atmosphere described with a diffusion equation. As refixation of respired and photorespired CO_2_ was shown to range between 24-38 % in wheat and rice [32], a static CO_2_ refixation of 25 % from mitochondiral glycine decarboxylation was assumed with the remaining CO_2_ lost to the atmosphere. For APs that relocate the CO_2_ to the chloroplast we assumed 100 % of this could potentially be refixed. Ammonia was also modelled as a dynamic variable along with ammonium assimilation into glutamate and the associated energy costs. The model was built in a stepwise manner, with each iteration being compared to the previous one to ensure that the changes made were valid (see supplementary material section S2 for a complete description).The final model was used as a reference point, called WT, and the APs were implemented on top of this model. Native photorespiration was deactivated in the presence of the APs by setting the Vmax of the first native photorespiratory enzyme that was not used by the respective AP to zero. The AP designs were grouped by their CO_2_ stoichiometry and the best performing pathway of each group is shown in the results. All code required to reproduce the results is available at https://gitlab.com/gain4crops/2024-paper.

## S1 Supplementary material

**Figure S1:**
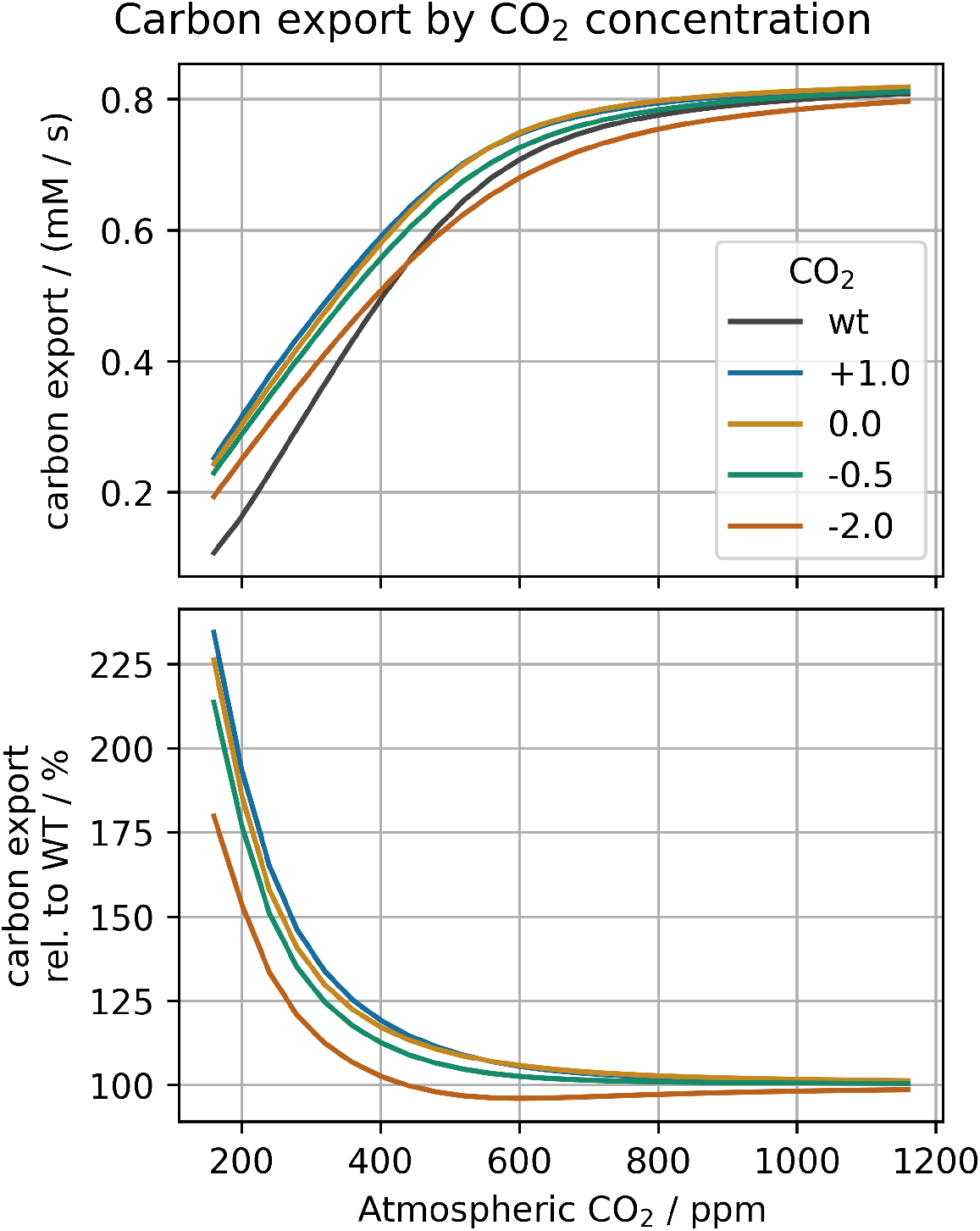
Steady-state carbon export of photorespiratory APs simulated by the kinetic model depending on atmospheric CO_2_ concentration in absolute units (top) and relative to WT (bottom). The AP designs are grouped by their CO_2_ stoichiometry and the maximal carbon export of the group is shown.

**Figure S2:**
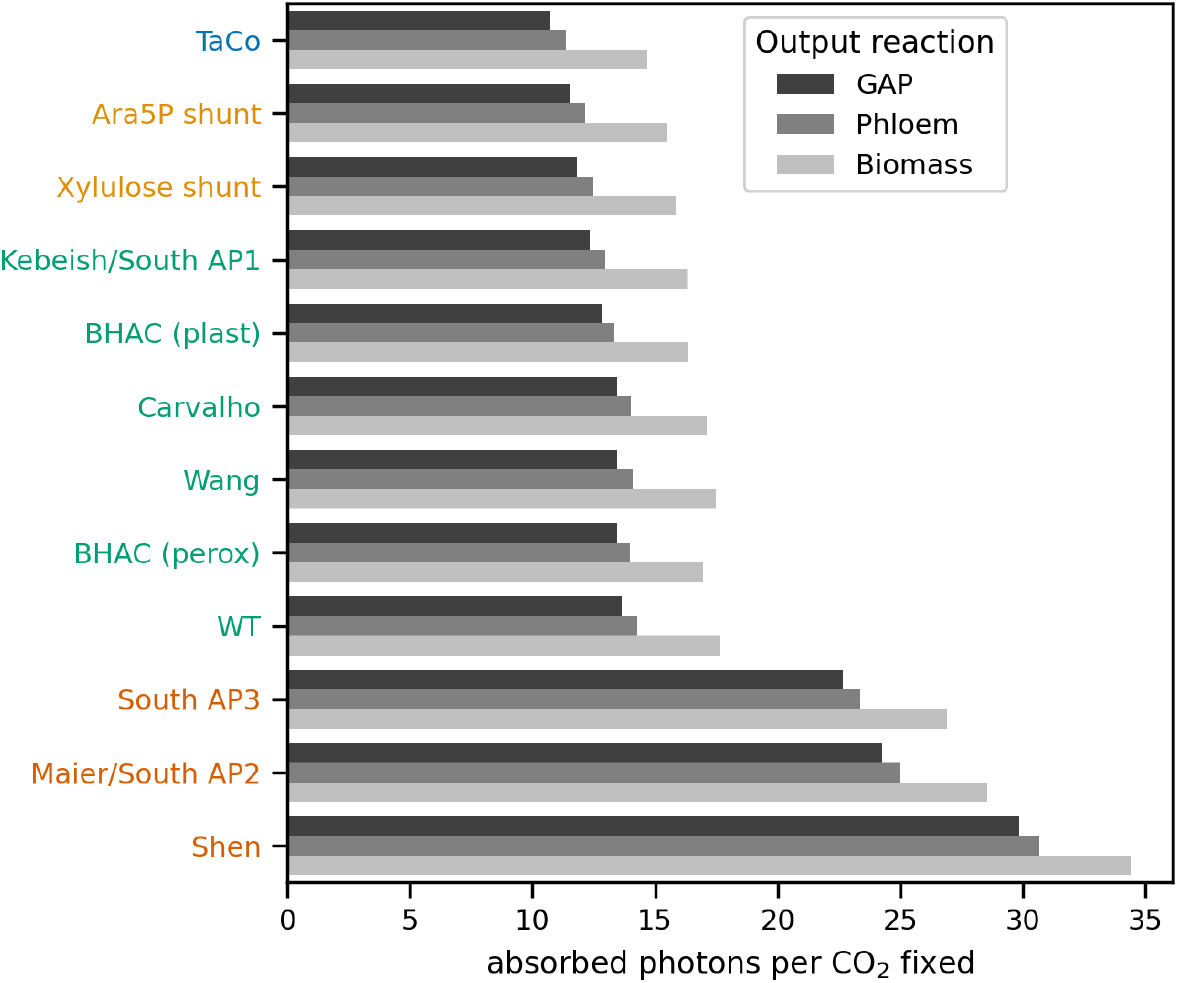
Absorbed photons per CO_2_ fixed for different APs with a rubisco rubisco carboxylase:oxygenase fixed at 3:1 calculated by stoichiometric FBA modelling. Model outputs were fixed to either glyceraldehyde 3-phosphate (GAP), phloem exudate or biomass represented by grey shading. AP label colours signify the amount of CO_2_ the AP fixed or released by the alternative pathways; blue, 1; yellow, 0; green, -0.5; orange, -2.

## S2 Kinetic model building

In the following, each model iteration is described in order to justify assumptions made. The analysis was started using the Poolman 2000 implementation of the Pettersson 1988 model. Besides the canonical reactions of the CBB cycle this model contains a saturating term to describe ATP generation but a fixed concentration of NADPH and NADP, oxygen and carbon-dioxide. Further, it contains a moiety for orthophosphate and another one to control the total efflux of the three triose-phosphates 3PGA, dihydroxyacetone phosphate (DHAP) and GAP, which besides the G1P efflux are the only exporter reactions. For a complete description, see the Pettersson 1988 and Poolman 2000 papers respectively [38, 49].

### Ambient CO2 conditions

The Pettersson and Ryde-Pettersson CBB cycle model, as well as the Poolman implementation, assume saturating CO2 conditions, which in turn also inhibits any photorespiratory activity. As the following analyses are all performed under ambient CO2 and O2 concentrations, the first change was to add a static internal CO2 concentration. We assumed an internal CO2 concentration of 0.012 mM, which corresponds to roughly 50 ppm - well in the range of 10-100 ppm CO2 measured in plant cells, see table S1.

Fig. S3 shows the relative difference in concentrations and fluxes between the model in saturating and ambient CO2 concentrations. As was found by Pettersson and Ryde-Pettersson, most fluxes are not very sensitive to the CO2 concentration in this range, with rubisco showing only a roughly 2.5 % decline in activity, even though the CO2 concentration was reduced to 6 % of its previous value. At the same time, the largest relative negative and positive changes are for 3PGA and RUBP respectively, which are product and substrate of the rubisco carboxylation. This suggests that the CBB cycle, or at least this model of the cycle, is buffered to keep the other intermediate concentrations stable, justifying further analysis and validity of the model at this changed CO2 concentration. For a deeper discussion of this behaviour see both Pettersson 1988 and Poolman 2000.

**Figure S3:**
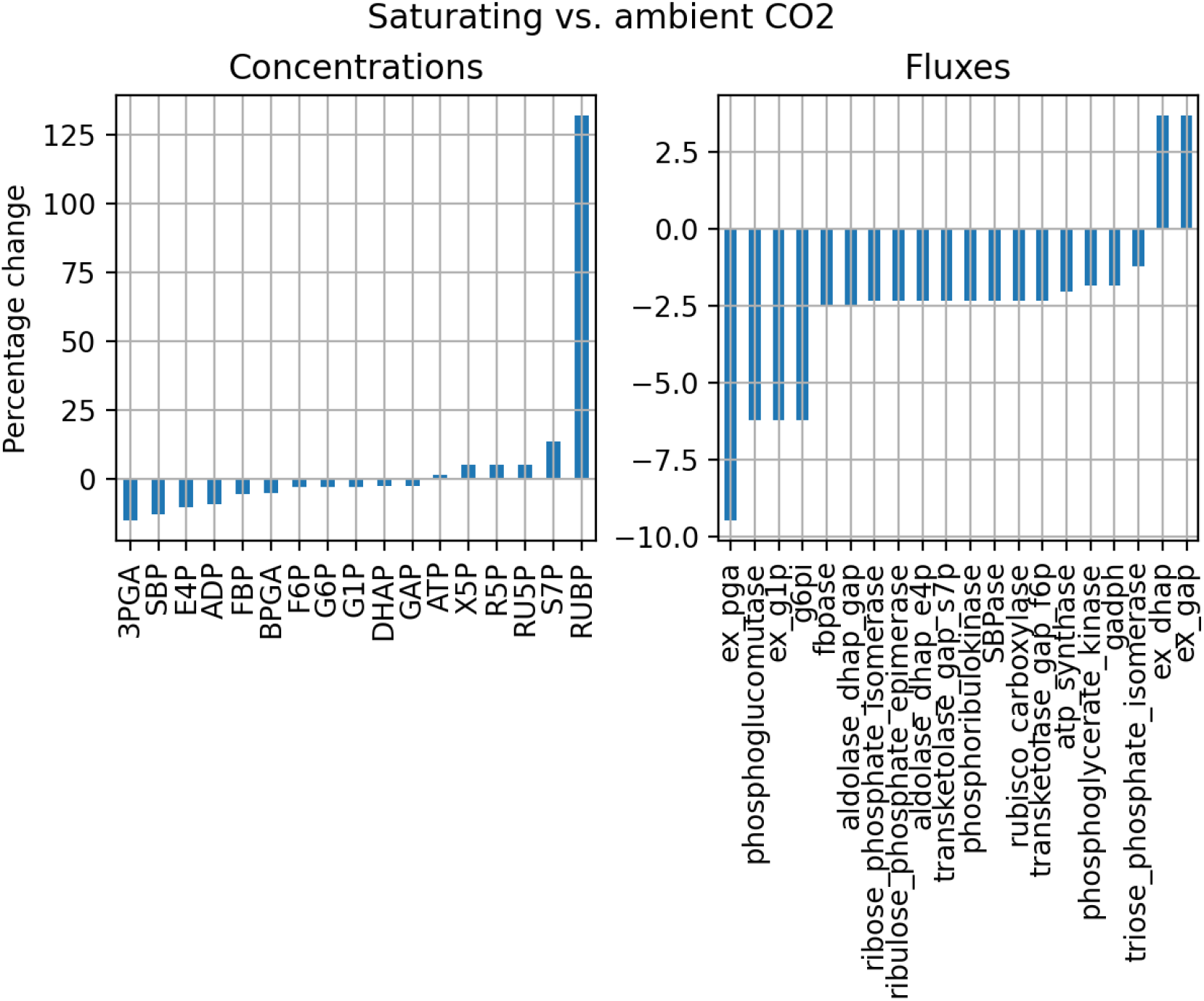
Comparison of saturating and ambient CO2 concentration. Shown on the are the relative differences between the concentrations and fluxes of the concentrations on the left and fluxes on the right.

#### S2.1.1 Poolman model at ambient CO2 changes

##### Parameters

**Table S1:** Parameters of Poolman model at ambient CO2

| Parameter name | Parameter value |
| --- | --- |
| CO2 | 1.20e-02 |

### Witzel rubisco kinetics

In the Poolman model the description of rubisco just contains the carboxylase reaction and no oxygenase activity. It is described using Michaelis-Menten kinetics with competitive inhibition terms by 3PGA, FBP, SBP, Pi and NADPH.

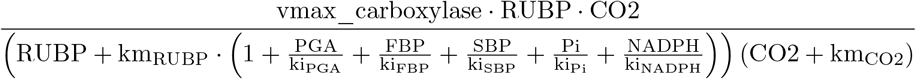

However, this description does not contain the complex interaction of both CO2 and O2 with rubisco. In the description by Witzel 2010, more emphasis is put on the interaction between CO2 and O2, while the competitive inhibition terms by Poolman are not included. The rate equation for the carboxylase is

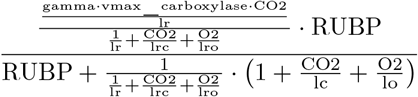

and for the oxygenase

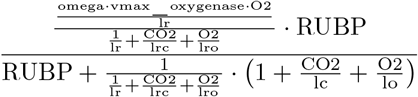

Note that we also did not include the inhibition terms by 3PGA, FBP, SBP, Pi and NADPH into the rate equations, but kept the original description by Witzel. The kinetic parameters of which can be found in table S3; for a full description of the rate equations and stoichiometries refer to section S2.2.2.

As 2PG also carries a phosphate group, the moiety function for total phosphate was changed as well, see section S2.2.2.

As at this point the Yokota 1985 model of photorespiration was not yet added, a temporary lumped photorespiration reaction was introduced to replenish the 3PGA otherwised lost. This reaction has the following stoichiometry: pgo + atp ⇋ 0.5 · pga and is described using mass-action kinetics:

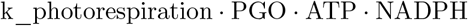

#### S2.2.1 Anoxic conditions

To first just asses the change of the rate equation to the system without also having to consider the complex dynamics of additional energy requirement by photorespiration we simulated the model in anoxic conditions. The expectation here is that this change in the description should have a minimal effect on the system, as long as the enzyme concentration is set appropriately.

As expected, in comparison to the model with the Pettersson rubisco kinetics at ambient CO2 concentration, Fig. S4 shows only minor changes in both concentrations and fluxes when the Witzel rubisco kinetics are used. This, again, justifies further analyses with this model iteration.

**Figure S4:**
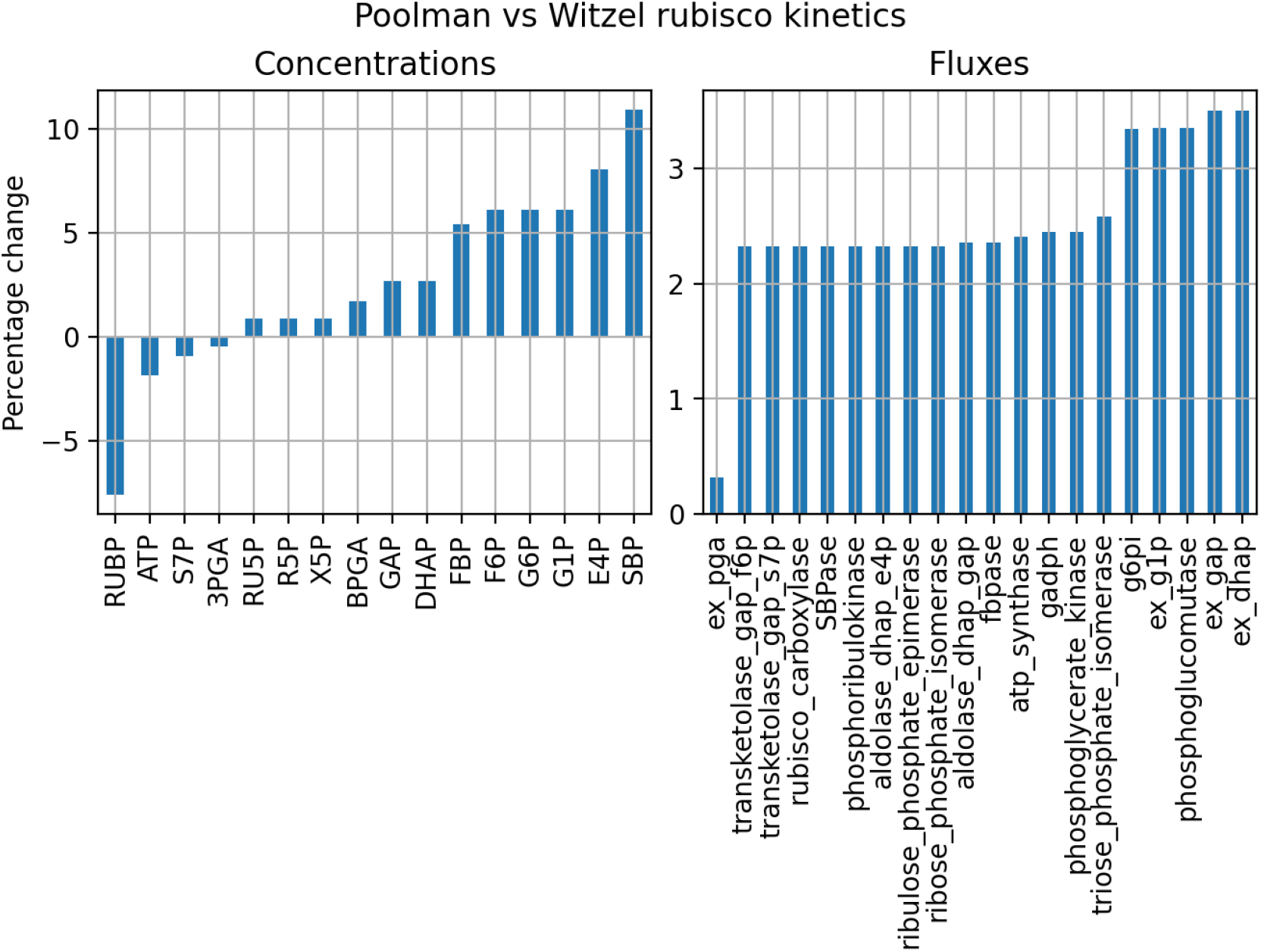
Comparison of rubisco kinetics by Pettersson 1988 and Witzel 2010. Shown on the are the relative differences between the concentrations and fluxes of the concentrations on the left and fluxes on the right.

#### S2.2.2 Witzel model at anoxia changes

##### Variables

**Table S2:** Variables of Witzel model at anoxia

| Model name | Name |
| --- | --- |
| pgo |  |

##### Parameters

**Table S3:** Parameters of Witzel model at anoxia

| Parameter name | Parameter value |
| --- | --- |
| E0_pgk | 1.00e+00 |
| E0_rubisco | 4.70e-01 |
| O2 | 0.00e+00 |
| gamma | 9.35e+01 |
| gamma_ker_plus | 1.40e+04 |
| k_er_minus | 4.80e-03 |
| k_er_plus | 1.50e+02 |
| k_pr | 1.00e+00 |
| kcat_pgk | 5.37e+02 |
| kcat_rcarb | 3.10e+00 |
| kcat_rox | 1.12e+00 |
| keq_pgk | 3.70e-04 |
| km_co2 | 1.07e-02 |
| km_o2 | 2.95e-01 |
| km_pgk_p | 2.70e-01 |
| km_pgk_s | 1.80e-01 |
| lc | 4.57e-04 |
| lo | 1.26e-03 |
| lr | 3.20e-05 |
| lrc | 3.42e-07 |
| lro | 9.44e-06 |
| omega | 3.39e+00 |
| omega_kcat_carb | 1.05e+01 |
| omega_ker_plus | 5.08e+02 |
| omega_koxy | 3.81e+00 |
| vmax_pgk | 5.37e+02 |
| vmax_rcarb | 1.46e+00 |
| vmax_rox | 5.29e-01 |

##### Derived Parameters

**Table S4:**
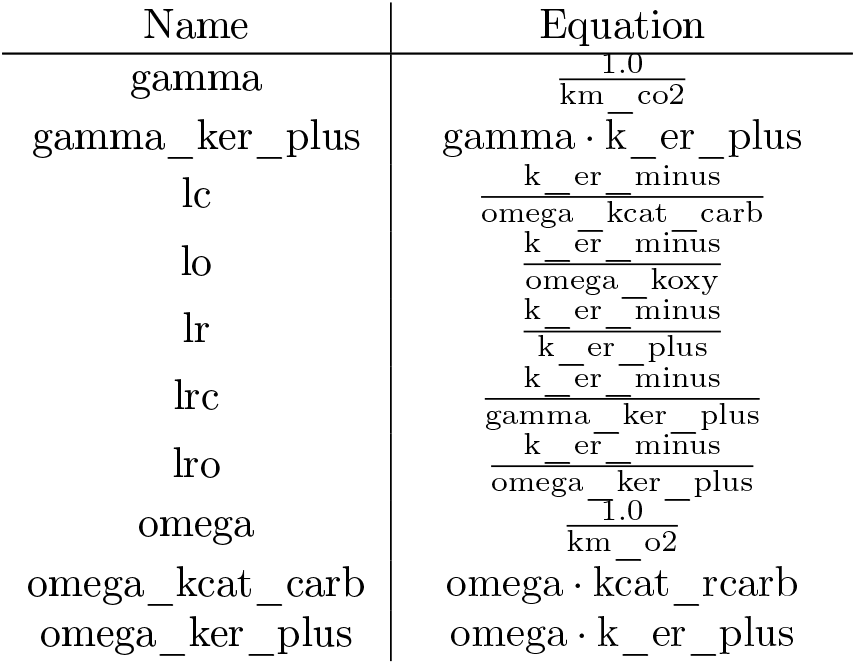
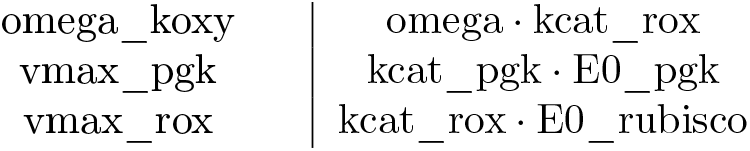
Derived parameters of Witzel model at anoxia

##### Algebraic Modules

###### pi

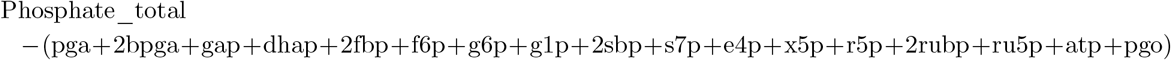

##### Reactions

###### phosphoglycerate kinase (PGK)

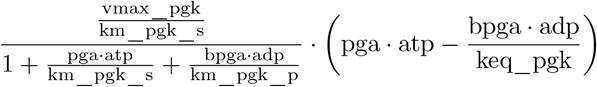

###### photorespiration (PR)

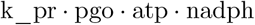

###### ribulose-1,5-bisphosphate carboxylase (RCARB)

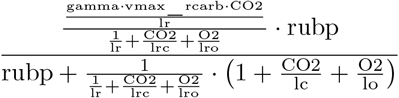

###### ribulose-1,5-bisphosphate oxygenase (ROX)

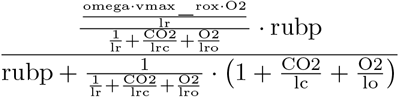

###### Stoichiometries

**Table S5:** Stoichiometries of Witzel model at anoxia

| Rate name | Stoichiometry |
| --- | --- |
| photorespiration (PR) | $\text{pgo} + \text{atp} \rightleftharpoons 0.5 \cdot \text{pga}$ |
| ribulose-1,5-bisphosphate oxygenase (ROX) | $\text{rubp} \rightleftharpoons \text{pga} + \text{pgo}$ |

#### S2.2.3 Ambient O2 conditions

The next step was to increase the oxygen concentration to ambient levels, which in turn led to production of 2-phosphoglycerate. The relative difference of concentrations and fluxes between anoxia and ambient oxygen levels are shown in Fig. S5. As the increased pull on the pool of RUBP when both oxygenase and carboxylase reactions of rubisco are active depleted the pool sizes of the CBB cycle intermediates, the export parameters for both triose-phosphate and G1P exporters were reduced, which is clearly visible in the decrease of 20-40 % in the respective fluxes. With these parameters, the 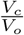 ratio was 3.65, which is within the range of experimentally determined values and thus the model predictions were deemed sufficient to continue the investigation.

**Figure S5:**
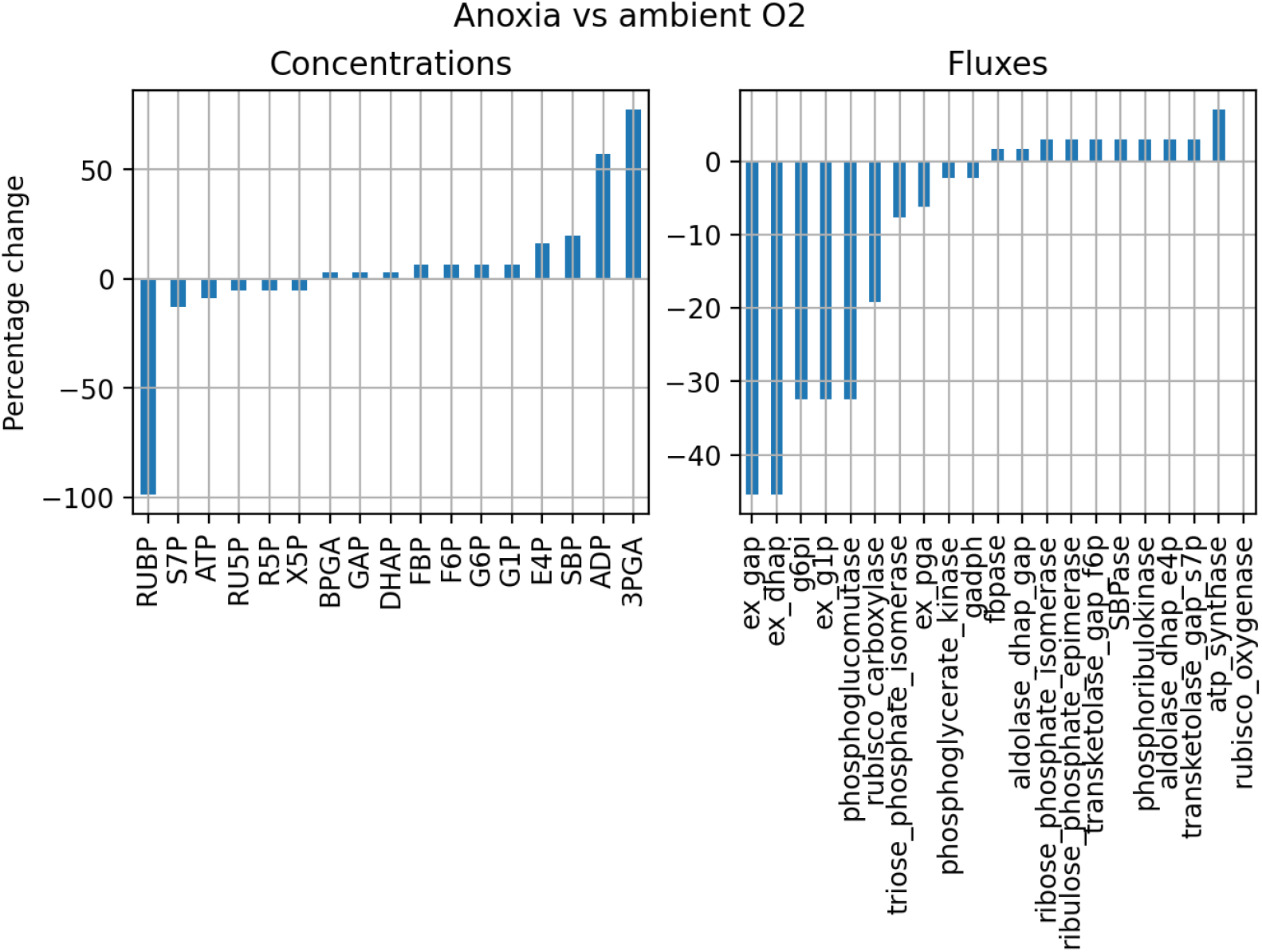
Comparison of anoxia and ambient O2. Shown on the are the relative differences between the concentrations and fluxes of the concentrations on the left and fluxes on the right.

The adjusted paramters can be found in table S6.

#### S2.2.4 Witzel model at ambient O2 changes

##### Parameters

**Table S6:** Parameters of Witzel model at ambient O2

| Parameter name | Parameter value |
| --- | --- |
| E0_ex_g1p | 5.00e-01 |
| E0_ex_tp | 6.00e-01 |
| E0_rubisco | 7.00e-01 |
| O2 | 2.50e-01 |
| k_pr | 1.00e+03 |
| vmax_ex_dhap | 1.20e+00 |
| vmax_ex_g1p | 1.60e-01 |
| vmax_ex_gap | 1.20e+00 |
| vmax_ex_pga | 1.20e+00 |
| vmax_rcarb | 2.17e+00 |
| vmax_rox | 7.87e-01 |

### Addition of Yokota photorespiration model

Next, we merged our model with the Yokota 1985 photorespiratory model [39]. For this addition eight new compounds had to be added to the model: H2O2, glycolate, glyoxylate, hydroxypyruvate, serine, glycine and glycerate, while glutamate and oxoglutarate were implemented as a parameter and derived parameter respectively. While the Yokota model features a phosphoglycolate influx, we replaced this by a phosphoglycolate phosphatase, just as we added a glycerate dehydrogenase and a glycerate kinase instead of the hydroxypyruvate efflux.

The stoichiometries and rate equations of the implemented reactions can be found in section S2.3.1.

With those additions the model was still incapable of metabolising the entire influx of phosphoglycolate, as some reactions were too slow. This is due to the Yokota model being calibrated to a constant influx not matching the oxygenase flux present in our model. We thus increased the enzyme concentrations as necessary to match the provided flux, the resulting parameters can be found in table S8.

Fig. S6 shows the differences in concentrations and fluxes between the lumped photorespiration term and the Yokota photorespiration model, both at ambient oxygen and carbon-dioxide concentrations. While there are noticable differences in the 2PG, RuBP and sedoheptulose-1,7-bisphosphate (SBP) concentrations, the differences in fluxes are generally small and mainly in which of the triose-phosphates is exported. This is due to the reduction in 3PGA concentration and increase in GAP and DHAP concentrations, leading to a decrease in the 3PGA export, while the export rates of DHAP and GAP are increased.

**Figure S6:**
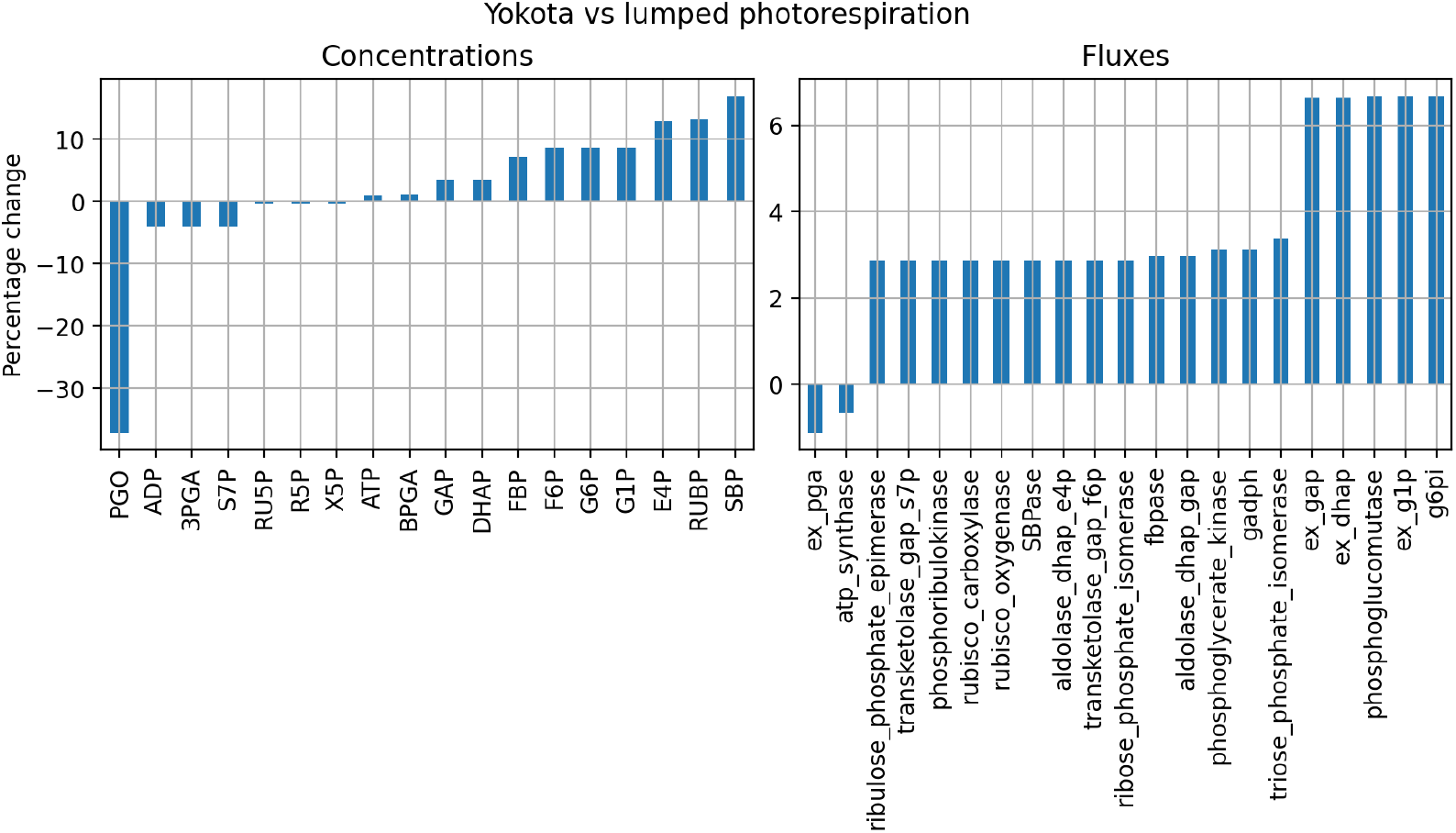
Comparison of lumped photorespiration and Yokota photorespiration model. Shown on the are the relative differences between the concentrations and fluxes of the concentrations on the left and fluxes on the right.

#### S2.3.1 Witzel-Yokota model changes

##### Variables

**Table S7:** Variables of Witzel-Yokota model

| Model name | Name |
| --- | --- |
| gly | glycine |
| glyc | glycerate |
| glyco | glycolate |
| glyx | glyoxylate |
| h2o2 | hydrogen peroxide |
| hpyr | hydroxypyruvate |
| ser | serine |

##### Parameters

**Table S8:** Parameters of Witzel-Yokota model

| Parameter name | Parameter value |
| --- | --- |
| E0_cat | 5.00e+01 |
| E0_gdc | 5.00e+01 |
| E0_gdh | 5.00e+01 |
| E0_gk | 5.00e+02 |
| E0_glcox | 1.00e+03 |
| E0_gta | 5.00e+01 |
| E0_pgp | 5.00e+01 |
| E0_sgta | 5.00e+00 |
| Glu_Oxo_total | 3.00e+00 |
| NAD_total | 8.60e-01 |
| glu | 2.00e+00 |
| kcat_cat | 2.11e+02 |
| kcat_gdc | 2.78e-02 |
| kcat_gdh | 1.11e-01 |
| kcat_gk | 1.59e-03 |
| kcat_glcox | 2.78e-02 |
| kcat_gta | 3.97e-02 |
| kcat_pgp | 8.11e-02 |
| kcat_sgta | 4.42e-02 |
| keq_gdh | 8.70e+04 |
| keq_gta | 3.00e+01 |
| keq_pgp | 3.10e+05 |
| keq_sgta | 6.00e+00 |
| ki_phosphoglycolate_phosphatase_Orthophosphate | 1.20e+01 |
| km_cat | 1.38e+02 |
| km_gdc_s | 6.00e+00 |
| km_gdh_p | 1.00e+00 |
| km_gdh_s | 1.20e-01 |
| km_glcox_s | 6.00e-02 |
| km_glycerate_kinase_glycerate | 2.50e-01 |
| km_gta_p | 1.00e+00 |
| km_gta_s | 3.00e+00 |
| km_pgp_p | 1.00e+00 |
| km_pgp_s | 2.90e-02 |
| km_sgta_p | 1.00e+00 |
| kms_transaminase_glyxoylate | 1.50e-01 |
| kms_transaminase_serine | 2.72e+00 |
| nad | 6.40e-01 |
| nadh | 2.20e-01 |
| nh4 | 1.00e+00 |
| oxogl | 1.00e+00 |
| vmax_cat | 1.06e+04 |
| vmax_gdc | 1.39e+00 |
| vmax_gdh | 5.53e+00 |
| vmax_gk | 7.94e-01 |
| vmax_glcox | 2.78e+01 |
| vmax_gta | 1.99e+00 |
| vmax_pgp | 4.06e+00 |
| vmax_sgta | 2.21e-01 |

##### Derived Parameters

**Table S9:** Derived parameters of Witzel-Yokota model

| Name | Equation |
| --- | --- |
| nad | $\text{NAD\_total} - \text{nadh}$ |
| oxogl | $\text{Glu\_Oxo\_total} - \text{glu}$ |
| vmax_cat | $\text{kcat\_cat} \cdot \text{E0\_cat}$ |
| vmax_gdc | $\text{kcat\_gdc} \cdot \text{E0\_gdc}$ |
| vmax_gdh | $\text{kcat\_gdh} \cdot \text{E0\_gdh}$ |
| vmax_gk | $\text{kcat\_gk} \cdot \text{E0\_gk}$ |
| vmax_glcox | $\text{kcat\_glcox} \cdot \text{E0\_glcox}$ |
| vmax_gta | $\text{kcat\_gta} \cdot \text{E0\_gta}$ |
| vmax_pgp | $\text{kcat\_pgp} \cdot \text{E0\_pgp}$ |
| vmax_sgta | $\text{kcat\_sgta} \cdot \text{E0\_sgta}$ |

##### Reactions

###### catalase (CAT)

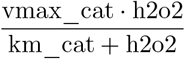

###### glycine decarboxylase (GDC)

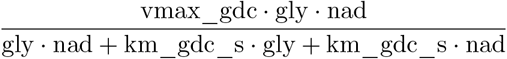

###### glutamate dehydrogenase (GDH)

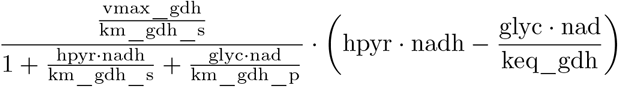

###### glycerate kinase (GK)

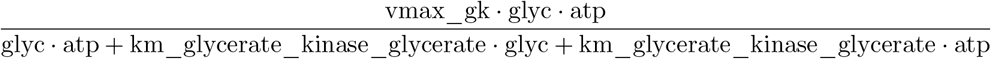

###### glycolate oxidase (GLCOX)

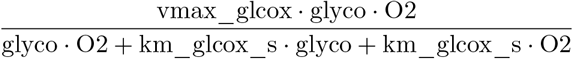

###### glycine transaminase (GTA)

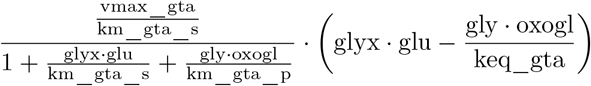

###### phosphoglycolate phosphatase (PGP)

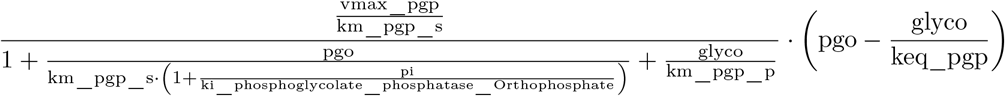

###### serine-glyoxylate transaminase (SGTA)

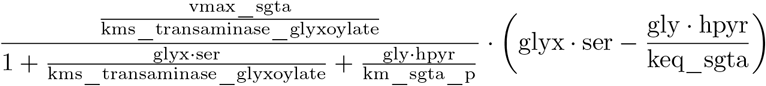

###### Stoichiometries

**Table S10:** Stoichiometries of Witzel-Yokota model

| Rate name | Stoichiometry |
| --- | --- |
| catalase (CAT) | h2o2 ⇌ ∅ |
| glycine decarboxylase (GDC) | 2.0 · gly ⇌ ser |
| glutamate dehydrogenase (GDH) | hpyr ⇌ glyc |
| glycerate kinase (GK) | glyc + atp ⇌ pga |
| glycolate oxidase (GLCOX) | glyco ⇌ glyx + h2o2 |
| glycine transaminase (GTA) | glyx ⇌ gly |
| phosphoglycolate phosphatase (PGP) | pgo ⇌ glyco |
| serine-glyoxylate transaminase (SGTA) | glyx + ser ⇌ gly + hpyr |

### Dynamic CO2

As the goal of this study is to compare photorespiratory bypasses, some of which decarboxylate 2-phosphoglycerate either once or twice, the effect that these additional decarboxylation reactions would have one the CO2 pool in the chloroplast was an important consideration. Thus, we decided to change the original description of a static CO2 concentration to a dynamic description, with a static external CO2 concentration at ambient levels (400 ppm) and a dissolving reaction. We described this dissolving reaction using diffusion kinetics

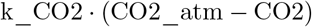

and fitted the rate constant to allow for the same flux as previously observed. As we wanted to keep the units consistent to mM, we used our previously assumed internal CO2 concentration of 50 ppm (=0.012 mM), which is 1/8 of the external CO2, and thus set the external CO2 concentration to be 0.012 · 8 = 0.096 mM. The parameters changed or added to the model can be found in table S12. Since CO2 was now a dynamic variable, the stoichiometries of all reactions consuming or producing CO2 needed to be adjusted, see table S2.4.1. Importantly, in order to model that the CO2 released by glycine decarboxylase (GDC) in the mitochondrium is not in fact all recovered, but some of which is lost, we adjusted the stoichiometry to account for that, by only recovering 25 % of CO2 released by GDC.

As expected, this change mostly affected the glyoxylate, glycine and serine pools as Fig. S7 shows, which are dependent on the flux of glycine decarboxylase.

**Figure S7:**
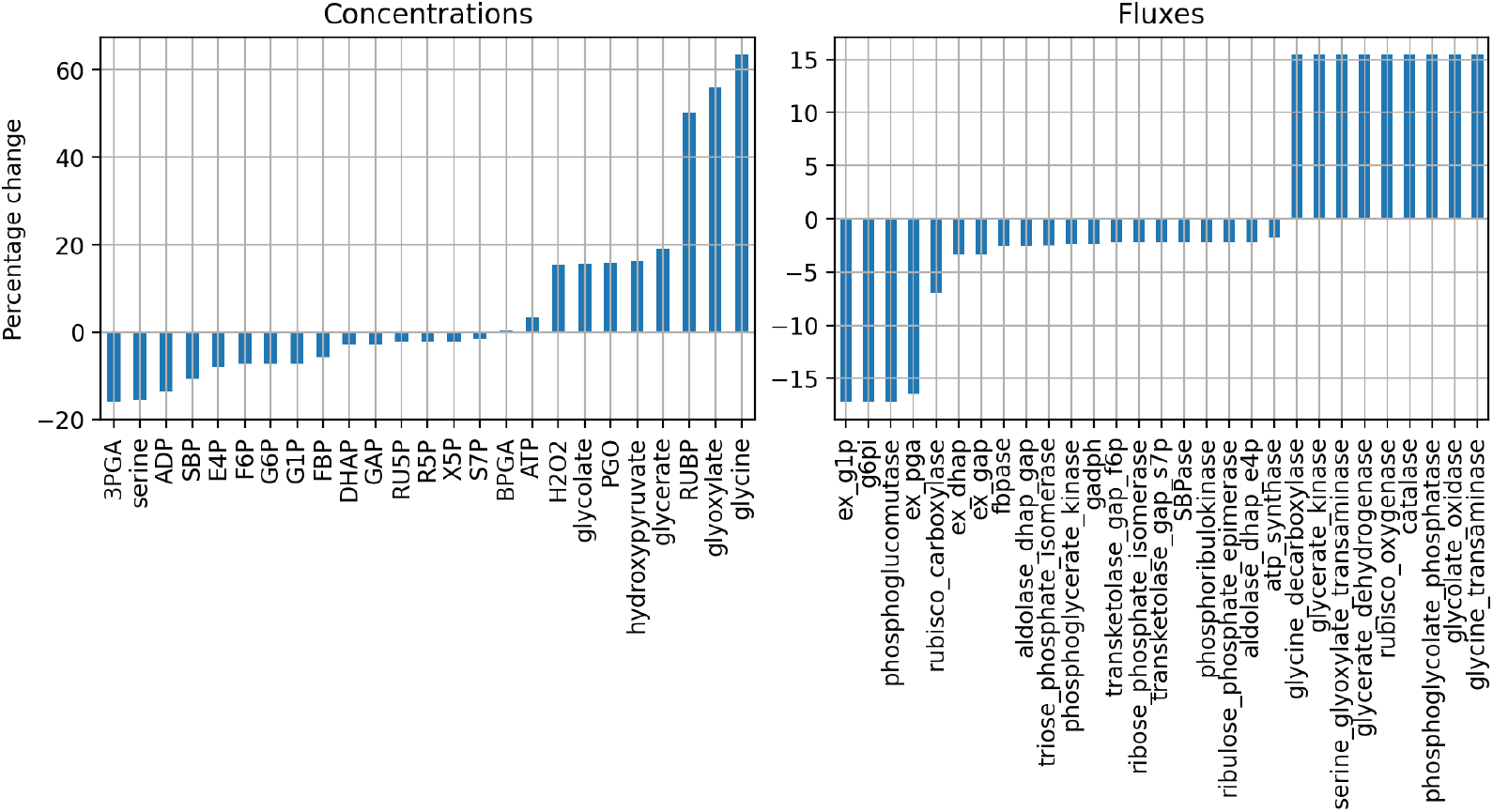
Comparison of dynamic and static CO2. Shown on the are the relative differences between the concentrations and fluxes of the concentrations on the left and fluxes on the right.

#### S2.4.1 dynamic CO2 model changes

##### Variables

**Table S11:** Variables of dynamic CO2 model

| Model name | Name |
| --- | --- |
| CO2 |  |

##### Parameters

**Table S12:**
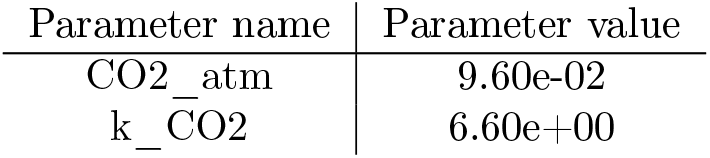
Parameters of dynamic CO2 model

##### Reactions

###### CO2 dissolving (CO2D)

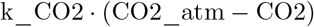

###### Stoichiometries

**Table S13:** Stoichiometries of dynamic CO2 model

| Rate name | Stoichiometry |
| --- | --- |
| CO2 dissolving (CO2D) | $\emptyset \rightleftharpoons CO2$ |
| glycine decarboxylase (GDC) | $2.0 \cdot gly \rightleftharpoons ser + 0.25 \cdot CO2$ |
| ribulose-1,5-bisphosphate carboxylase (RCARB) | $rubp + CO2 \rightleftharpoons 2.0 \cdot pga$ |

### Dynamic NADPH

Some of the alternative pathways use NAD(P)H / NAD(P) as redox pairs, while others use O2 / H2O2. In order to account for the difference in usable reduction equivalents, the static description of NADPH in the Pettersson model is insufficient. We thus implemented NADPH as a dynamic variable, NADP as an algebraic quantity derived from the NADPH concentration and a total pool size, and a ferredoxin-NADP^+^ reductase (FNR) that generates NADPH from NADP. This description follows the one by Matuszyńska 2019 [51]. The kinetic parameters, stoichiometries and rate equations for this model iteration can be found in section S2.5.1. Fig. S8 shows the relative changes in concentrations and fluxes caused by this change between a static and dynamic description of NADPH. While the fluxes are relatively similar, a noticable decrease in the concentrations of 2,3-bisphosphoglycerate (BPGA) and GAP can be found, while RuBP and SBP increase. This shows, that the static NADPH assumed by Pettersson et. al. might only be valid in very specific conditions, as the time course in Fig. S9 clearly shows, that the NADPH concentration changes rapidly to a new steady state from the previously assumed one.

**Figure S8:**
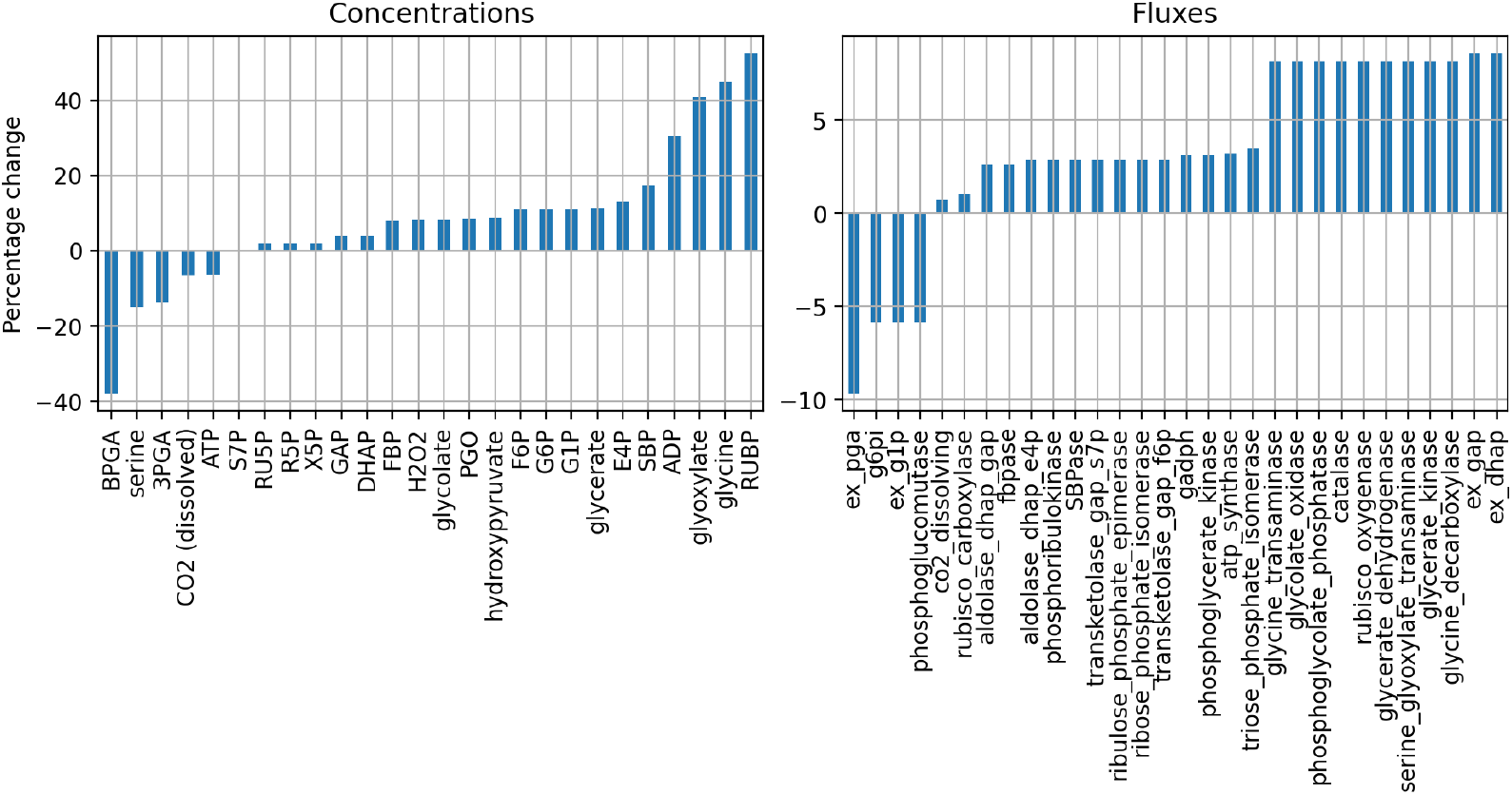
Comparison of static and dynamic NADPH. Shown on the are the relative differences between the concentrations and fluxes of the concentrations on the left and fluxes on the right.

**Figure S9:**
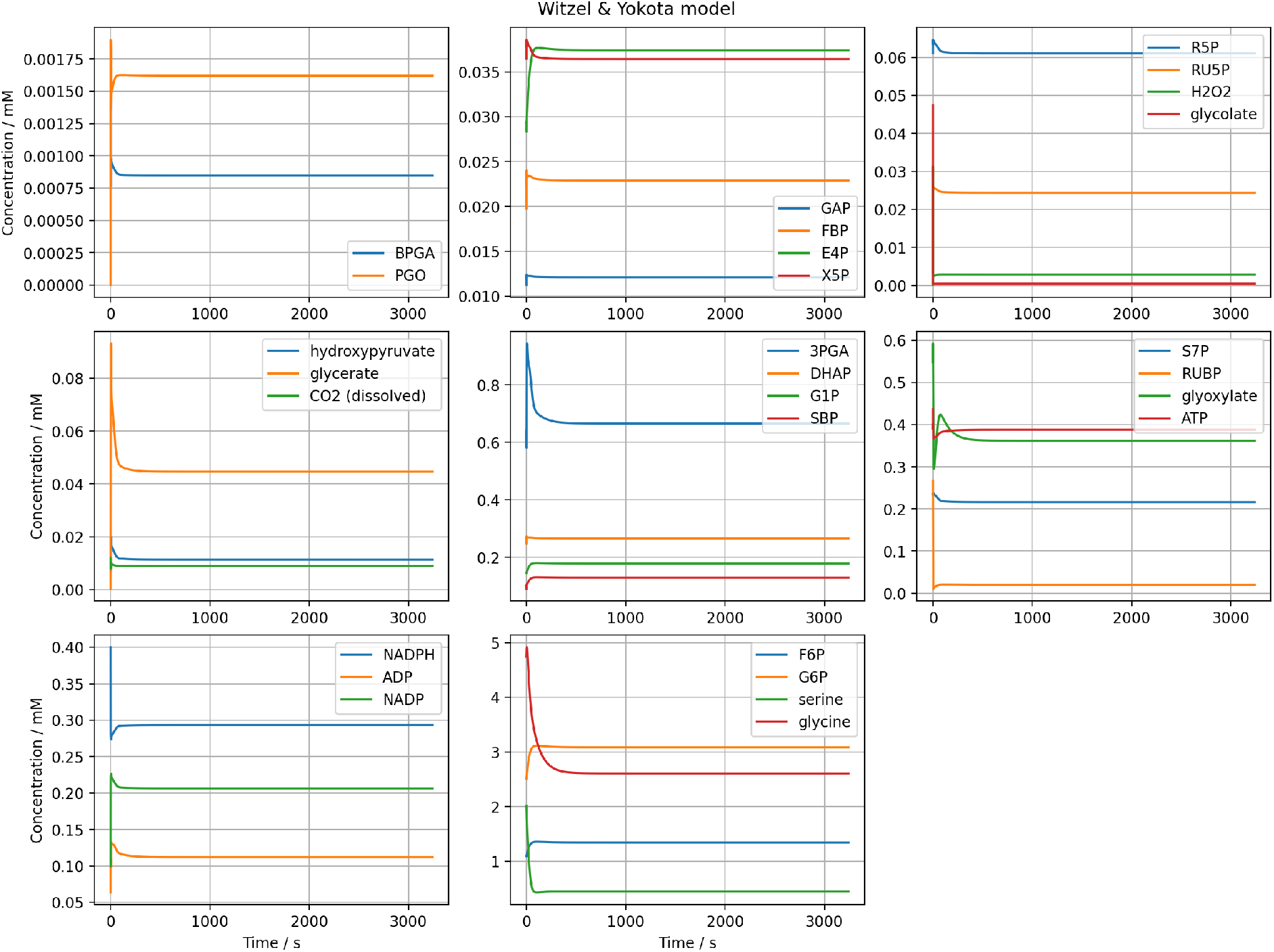
Time course of metabolite concentrations in model with dynamic NADPH

#### S2.5.1 dynamic NADPH model changes

##### Variables

**Table S14:** Variables of dynamic NADPH model

| Model name | Name |
| --- | --- |
| nadph | Nicotinamide adenine dinucleotide phosphate (reduced) |

##### Parameters

**Table S15:** Parameters of dynamic NADPH model

| Parameter name | Parameter value |
| --- | --- |
| E0_fnr | 1.00e+00 |
| kcat_fnr | 2.82e+00 |
| km_fnr | 1.90e-01 |
| vmax_fnr | 2.82e+00 |

##### Derived Parameters

**Table S16:** Derived parameters of dynamic NADPH model

| Name | Equation |
| --- | --- |
| vmax_fnr | kcat_fnr · E0_fnr |

##### Algebraic Modules

###### nadp

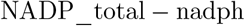

##### Reactions

###### ferredoxin:nadph reductase (FNR)

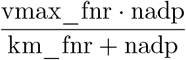

###### Stoichiometries

**Table S17:** Stoichiometries of dynamic NADPH model

| Rate name | Stoichiometry |
| --- | --- |
| ferredoxin:nadph reductase (FNR) | $\emptyset \rightleftharpoons \text{nadph}$ |
| glyceraldehyde 3-phosphate dehydrogenase (GADPH) | $\text{nadph} + \text{bpga} \rightleftharpoons \text{gap}$ |
| glycine decarboxylase (GDC) | $2.0 \cdot \text{gly} \rightleftharpoons \text{ser} + \text{nadph} + 0.25 \cdot \text{CO2}$ |
| glutamate dehydrogenase (GDH) | $\text{nadph} + \text{hpyr} \rightleftharpoons \text{glyc}$ |

### Thioredoxin regulation

Saadat 2021 found that a dynamic regulation of the CBB cycle activity based on the available energy is necessary to stabilise the system in a wide variety of light conditions [52]. Plants use the thioredoxin system to dynamically activate and deactivate rubisco, FBPase, SBPase, phosphoribulokinase and glycine decarboxylase based on the redox-state of the photosynthetic electron transport chain, specifically ferredoxin. As this model does not contain the PETC, and thus no ferredoxin concentration, we choose to implement an NADPH-thioredoxin reductase instead of the ferredoxin-thioredoxin reductase, but otherwise follow the implementation by Saadat 2021, which means that two new variables were added: the oxidised thioredoxin concentration as well as the fraction of CBB cycle enzymes that is active. The reduced thioredoxin concentration and inactive fraction of CBB cycle enzymes was expressed using conservation relations. This means that for rubisco, FBPase, SBPase, phosphoribulokinase and glycine decarboxylase the enzyme concentration now was described using kinetics like the following (with E0_rubisco_ being replaced by the respective enzyme concentration) E_active · E0_rubisco_.

Refer to section S2.6.1 for exact descriptions of this implementation.

As this change speeds up the CBB cycle, including rubisco, it was necessary to adjust some parameters like the rubisco concentration and maximal triose-phosphate export rates to stay within the range of experimentally observable concentrations. The kinetic parameters for this model iteration can be found in table S19. Fig. S10 shows the relative changes in concentrations and fluxes caused by the changes in this model iteration. As with most of the previous changes, the main differences can be seen in the triose-phosphate and glucose export rates, and their respective concentrations, as well as the balance between glycine and serine.

**Figure S10:**
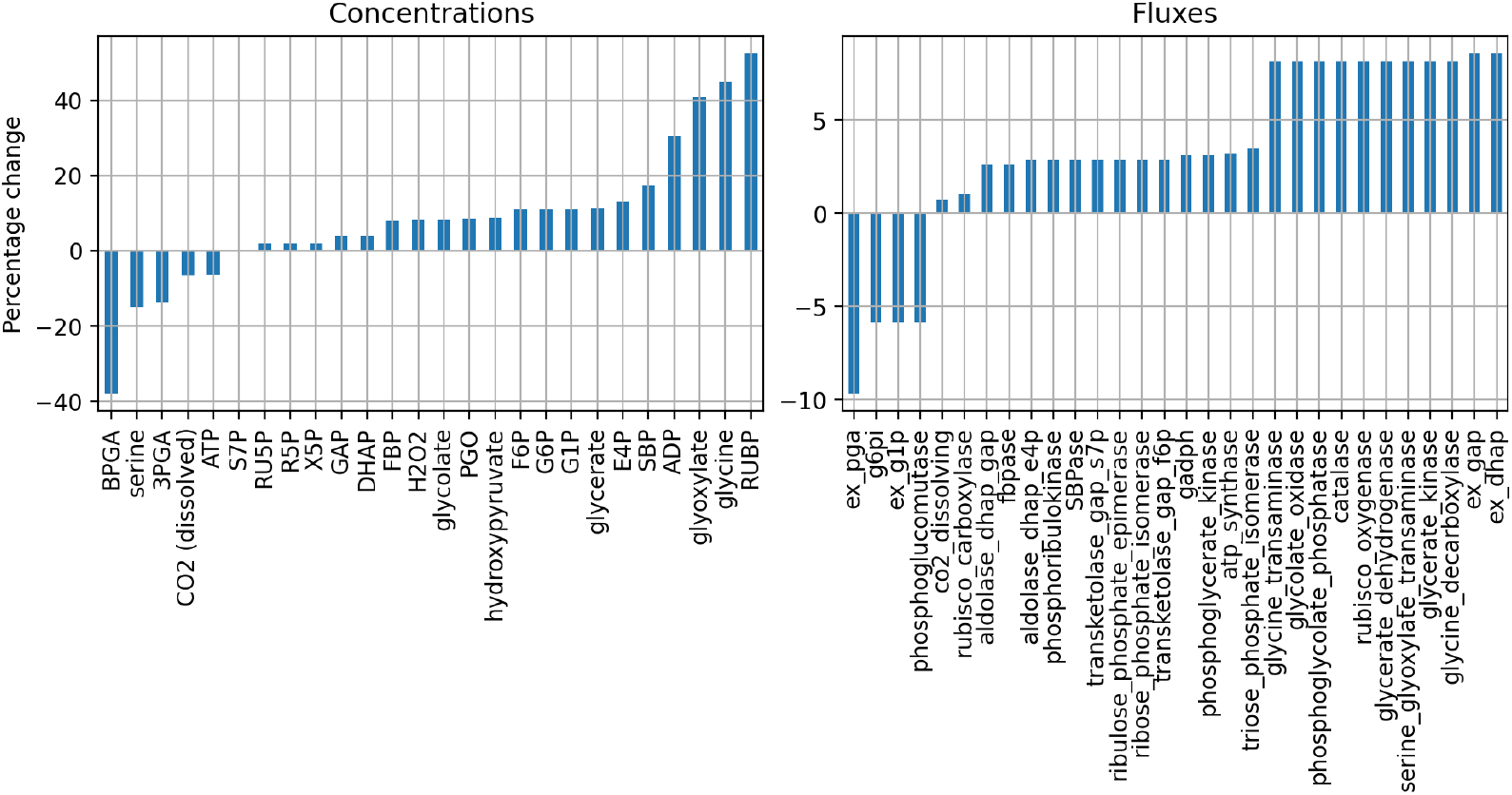
Comparison of thioredoxin-regulated and unregulated enzyme concentrations. Shown on the are the relative differences between the concentrations and fluxes of the concentrations on the left and fluxes on the right.

While the relative change in the concentrations of serine and glycine seems large, Fig. S11 clearly shows that both concentrations are predicted to be in acceptable magnitudes.

**Figure S11:**
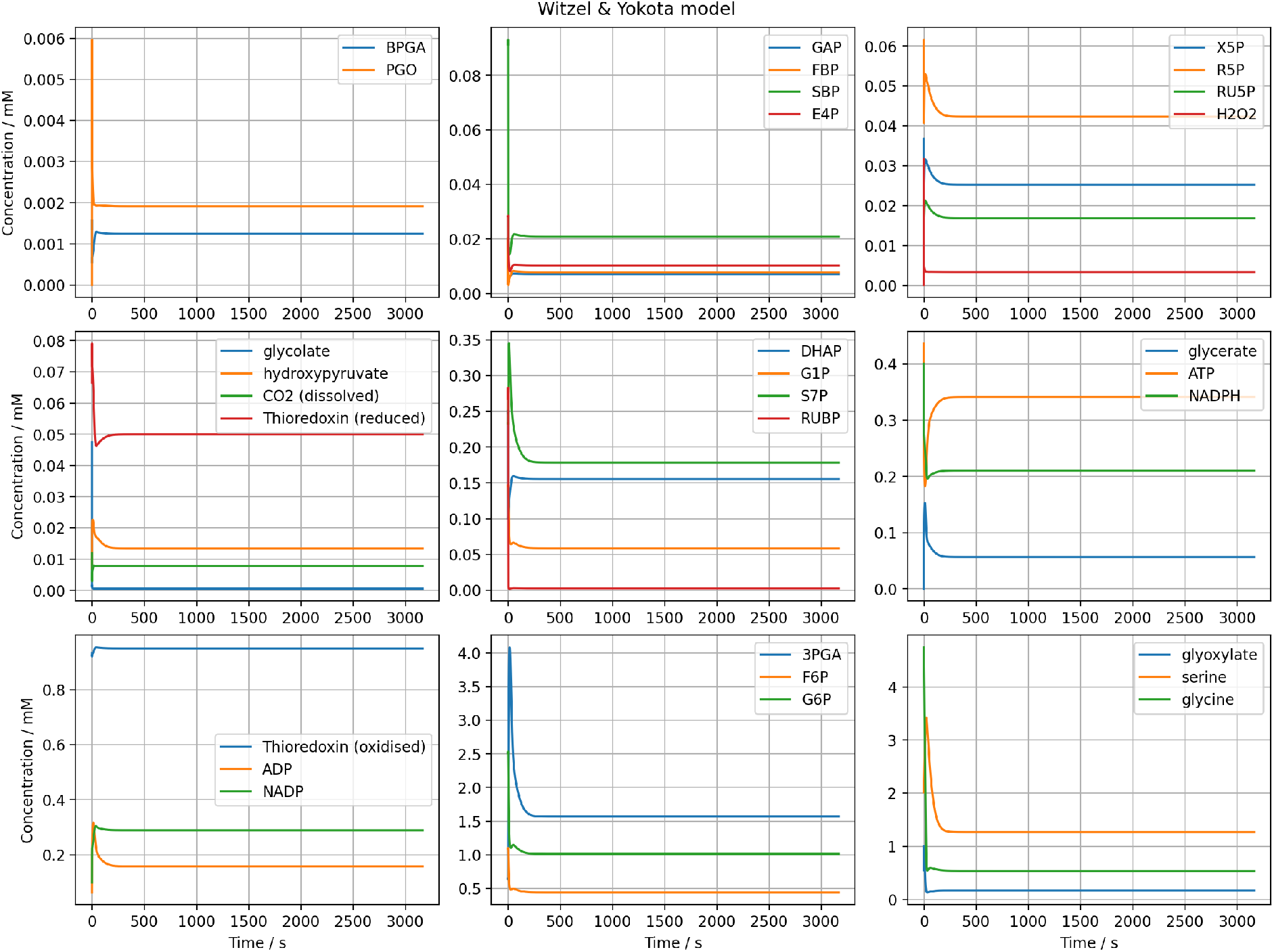
Time course of metabolite concentrations in model with thioredoxin regulation

#### S2.6.1 thioredoxin-regulated model changes

##### Variables

**Table S18:**
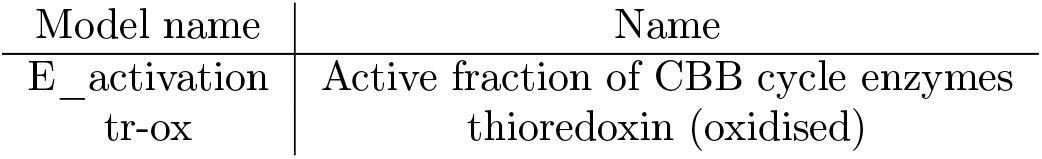
Variables of thioredoxin-regulated model

##### Parameters

**Table S19:** Parameters of thioredoxin-regulated model

| Parameter name | Parameter value |
| --- | --- |
| e_cbb_tot | 6.00e+00 |
| k_e_cbb_activation | 1.00e+00 |
| k_e_cbb_relaxation | 1.00e-01 |
| k_nadph_tr_reductase | 1.00e+00 |
| thioredoxin_tot | 1.00e+00 |

##### Algebraic Modules

###### E_fbpase

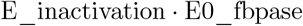

###### E_gdc

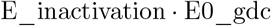

###### E_inactivation

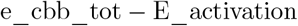

###### E_prk

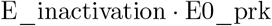

###### E_rubisco

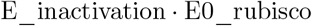

###### E_sbpase

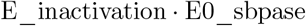

###### tr_red

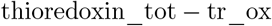

###### vmax_fbpase

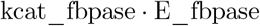

###### vmax_gdc

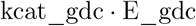

###### vmax_prk

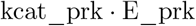

###### vmax_rcarb

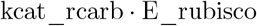

###### vmax_rox

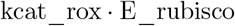

###### vmax_sbpase

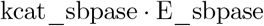

##### Reactions

###### Thioredoxin activation (E-ACT)

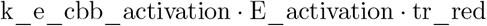

###### Thioredoxin deactivation (E-DEACT)

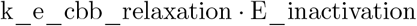

###### NADPH thioredixon reductase (NTR)

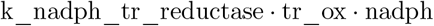

###### Stoichiometries

**Table S20:**
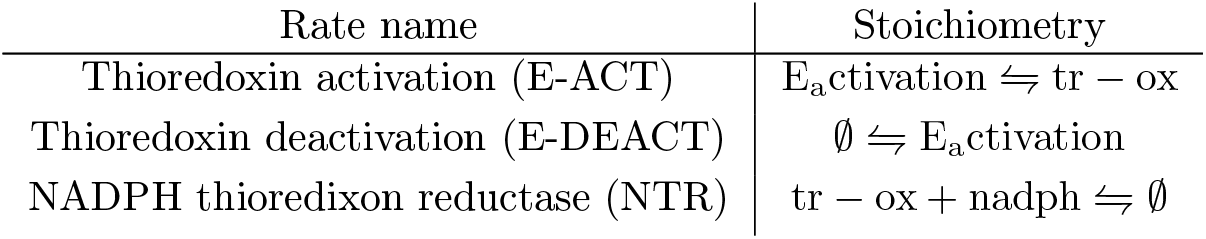
Stoichiometries of thioredoxin-regulated model

### Dynamic energy

With the previous model iteration, there were two independent maximal velocities for the production and NADPH and ATP that had no common source, which made it unintuitive to model energy availability due to illumination. In order to describe this behaviour in the simplest possible way we introduced a new ’energy’ variable, which is produced in a light-dependent manner and then used by both ATP synthase (ATPSYN) as well as ferredoxin:nadph reductase (FNR). This way, if the ratio between ATP and NADPH changes due to different sink requirements (e.g. different alternative pathways), the system can compensate for that, mimicking the role of cyclic electron flow. In order to also be able to estimate the amount of excess energy that could occur due to excessive illumination, we added an additional quenching reaction. Section S2.7.1 shows the exact implementations described before.

Fig. S12 shows the relative differences in concentrations and fluxes caused by the changes in this model iteration at ambient CO2 and O2 concentrations and a medium illumination of 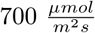.

**Figure S12:**
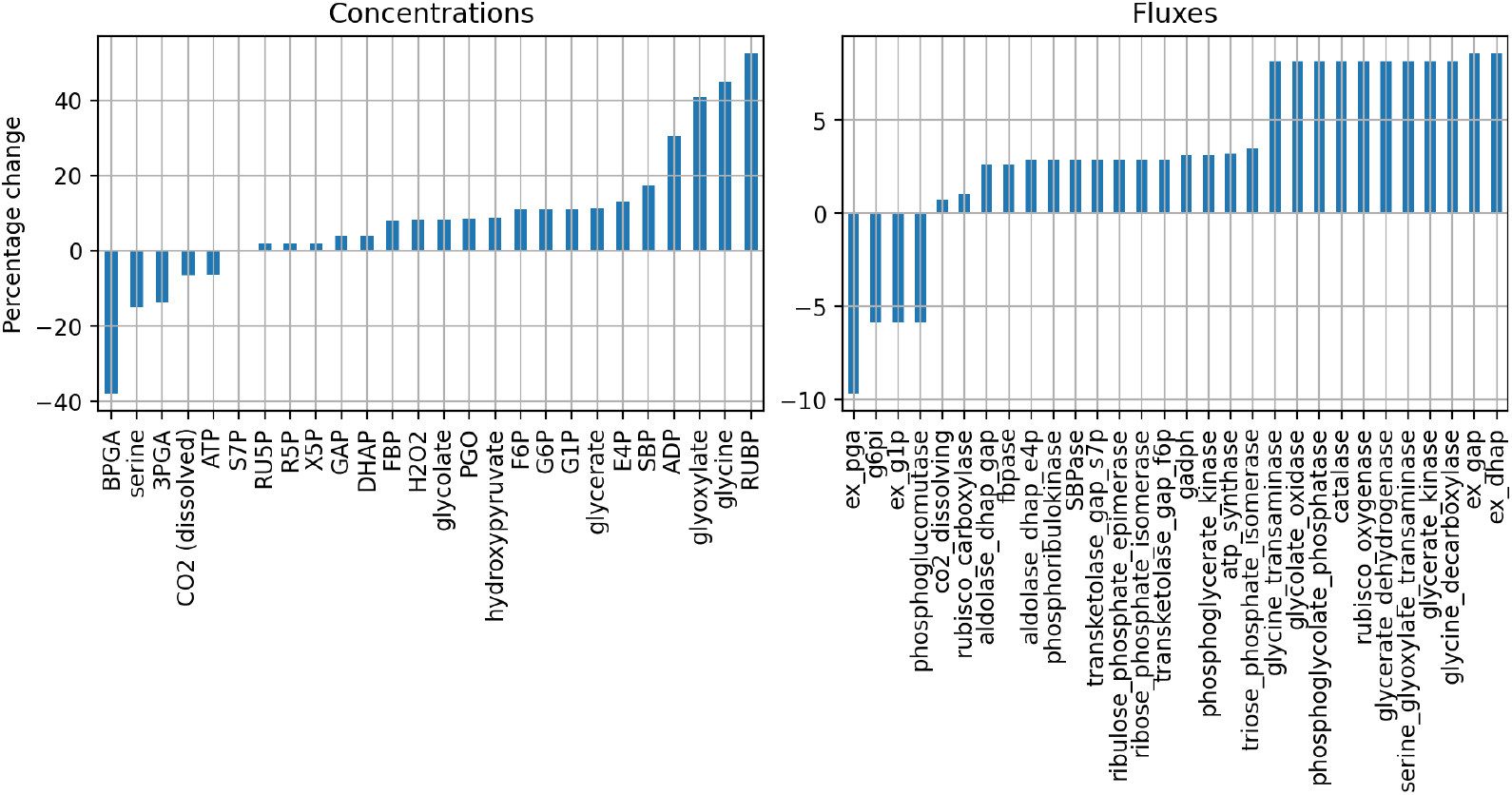
Comparison of static and dynamic NADPH. Shown on the are the relative differences between the concentrations and fluxes of the concentrations on the left and fluxes on the right.

#### S2.7.1 PPFD model changes

##### Variables

**Table S21:** Variables of PPFD model

| Model name | Name |
| --- | --- |
| energy |  |

##### Parameters

**Table S22:**
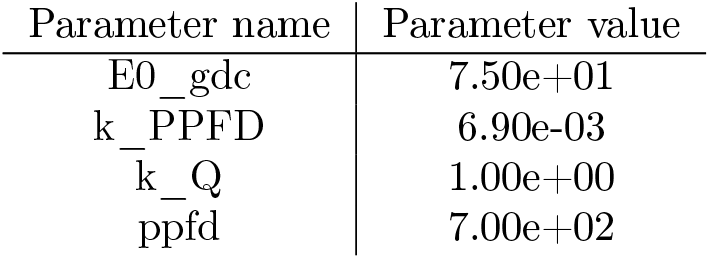
Parameters of PPFD model

##### Reactions

###### ATP synthase (ATPSYN)

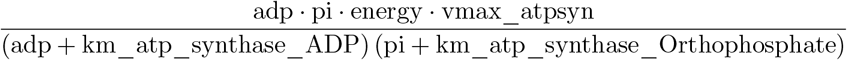

###### ferredoxin:nadph reductase (FNR)

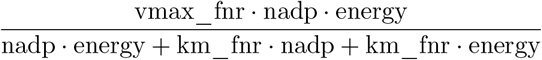

###### Photosynthetic electron transport chain (PETC)

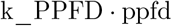

###### Quencher (Q)

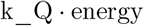

###### Stoichiometries

**Table S23:** Stoichiometries of PPFD model

| Rate name | Stoichiometry |
| --- | --- |
| ATP synthase (ATPSYN) | $\text{energy} \rightleftharpoons \text{atp}$ |
| ferredoxin:nadph reductase (FNR) | $\text{energy} \rightleftharpoons \text{nadph}$ |
| Photosynthetic electron transport chain (PETC) | $\emptyset \rightleftharpoons \text{energy}$ |
| Quencher (Q) | $\text{energy} \rightleftharpoons \emptyset$ |

### Dynamic ammonia

One of the advantages that some alternative pathways have over wild-type photorespiration is that they don’t required refixation of ammonia lost by glycine decarboxylase. For example, the BHAC pathway can either rely on spontaneous formation of oxaloacetate by deamination of iminoaspartate or use an aspartate:NADP+ oxidoreductase and an aspartate aminotransferase to convert iminoaspartate to aspartate and then oxaloacetate, while transferring the amino-group to alpha-ketoglutarate to form glutamate. This avoids the energetic cost of either glutamate dehydrogenase, which uses one NADPH to aminate oxoglutarate (OXOGL) in order to form glutamate (GLU), or using glutamine synthase (GS) and glutamine oxoglutarate aminotransferase (GOGAT), which use one ATP and one NADPH to refix the ammonia. We changed the previous static ammonia description to a dynamic one, to allow comparative analyses of these energetic differences, see section S2.8.1 for the exact implementation.

Fig. S13 shows the relative differences in concentrations and fluxes caused by the changes in this model iteration. The largest differences can be observed in the concentrations of glyoxylate and serine, which is likely caused by the difference in thermodynamic driving force of the now dynamic ammonium for GDC.

**Figure S13:**
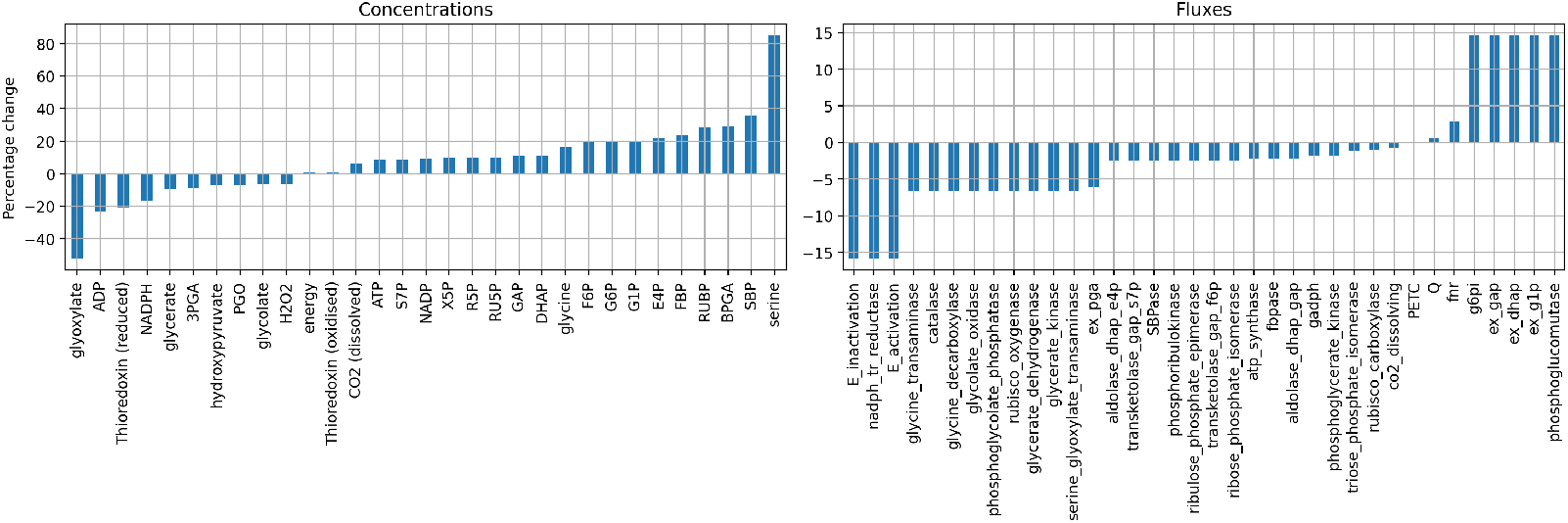
Comparison of static and dynamic NADPH. Shown on the are the relative differences between the concentrations and fluxes of the concentrations on the left and fluxes on the right.

#### S2.8.1 dynamic ammonium model changes

##### Variables

**Table S24:** Variables of dynamic ammonium model

| Model name | Name |
| --- | --- |
| glu | glutamate |
| nh4 | ammonium |

##### Parameters

**Table S25:** Parameters of dynamic ammonium model

| Parameter name | Parameter value |
| --- | --- |
| E0_gdh | 1.00e+00 |
| kcat_gdh | 1.04e+02 |
| keq_gdh | 7.20e+05 |
| km_gdh_p | 6.40e-01 |
| km_gdh_s | 1.54e+00 |
| vmax_gdh | 1.04e+02 |

##### Derived Parameters

**Table S26:**
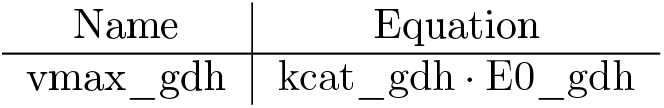
Derived parameters of dynamic ammonium model

##### Algebraic Modules

##### oxogl

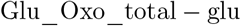

##### Reactions

###### glutamate dehydrogenase (GDH)

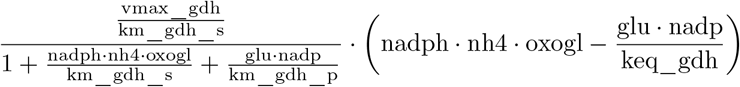

###### Stoichiometries

**Table S27:**
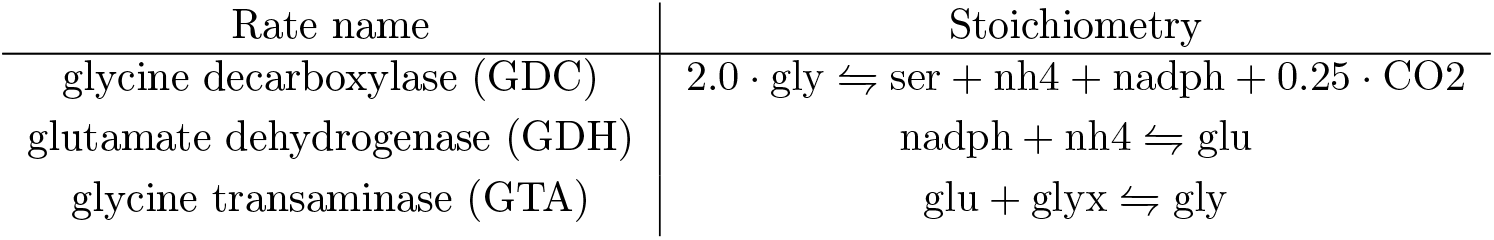
Stoichiometries of dynamic ammonium model

## S3 Analysis of wildtype model

Before adding the BHAC cycle, we wanted to make sure that the model we from now on refer to as ’wildtype’ behaves like expected. For this we systematically scanned the effect of illumination and external CO2 concentration on both rubisco carboxylase as well as carbon export, which we define as a weighted sum of the triose-phosphate exporters and the glucose exporters, to measure how many carbons are exported from the system:

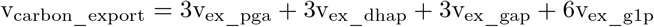

Fig. S14a shows the effect of different illumination of those selected rates. We can observe that both the ribulose-1,5-bisphosphate carboxylase (RCARB) and carbon export rate saturate with sufficiently high illumination. However, while the two rates are virtually identical at low illumination, the rubisco rate accelerates quicker than the carbon export rate. This suggests, that at high illumination the CBB cycle turns at a more rapid pace, but fewer carbons are being exported per revolution. A possible explanation for this is that at lower light the system is not bound by the diffusion of CO2 into the chloroplast, such that the internal CO2 concentration increases, leading to less oxygenation and a higher rubisco carboxylase:oxygenase. In that scenario the CBB cycle can spare some additional carbons, such that the carbon export rate approaches the carboxylase rate. On the other hand, in the high light scenario internal CO2 concentration becomes the limiting factor and the CBB cycle needs to retain more of it’s carbons to keep running, thus the carbon export rate can only be a fraction of the carboxylase rate.

**Figure S14:**
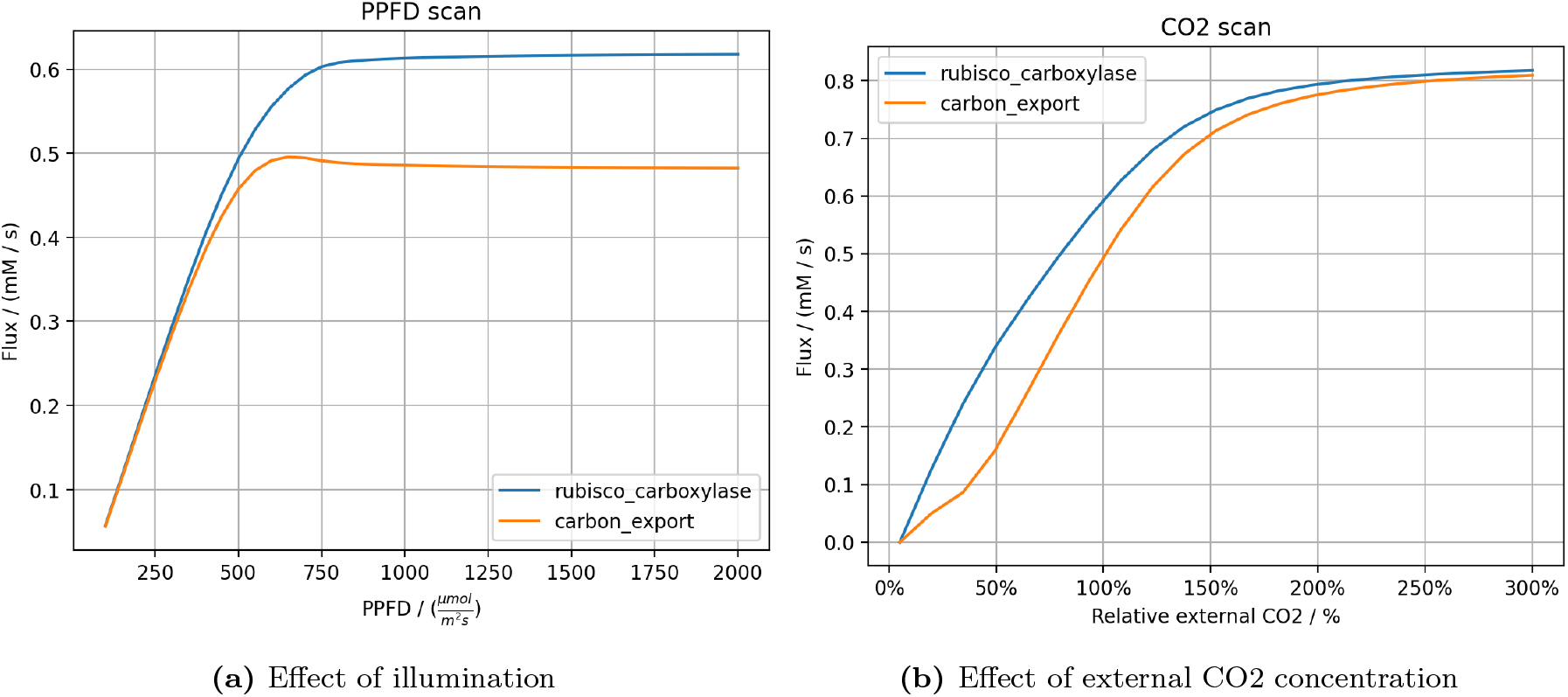
Systematic scans of illumination and external CO2 concentration on the rate of rubisco carboxylase and the rate of total carbon export.

This is supported by Fig. S14b, in which the external CO2 concentration was varied systematically. While at low external CO2 concentrations the fraction of carbon export to rubisco carboxylase is lower, at high external CO2 concentrations the carbon export again approaches the carboxylase rate. This analysis showed us that the model behaved as expected and already predicts a valuable strategy for photorespiratory bypasses: if the compartment of decarboxylation is moved from the mitochondrium to the chloroplast, then the internal CO2 concentration can be increased, as only a part of the CO2 released in the mitochondrium will diffuse into the chloroplast. This in turn can lead to a higher carbon export and thus better plant growth, as the oxygenase reaction is inhibited and more of the available CO2 can be used to be exported out of the chloroplast instead of using it to re-fuel the CBB cycle.

### Addition of the BHAC cycle

In the BHAC alternative pathway, glyoxylate is condensed with glycine to form hydroxyaspartate, which is dehydrated to iminoaspartate and spontaneously deaminated to oxaloacetate. Alternatively, a more efficient version reduces iminoaspartate to aspartate, which avoids ammonia release and thus saves energy. Asparate is then converted into oxaloacetate as well using a asparate aminotransferase. To replenish the CBB cycle, oxaloacetate is converted to malate, which in turn is decarboxylated to pyruvate. Pyruvate is then phosphorylated to phosphoenolpyruvate (PEP), cleaved by enolase to form and finally phosphoglycerate mutase isomerises 2PG into 3PGA. See Fig. S15 for an illustration and sectipm S3.1.1 for the reaction stoichiometries.

**Figure S15:**
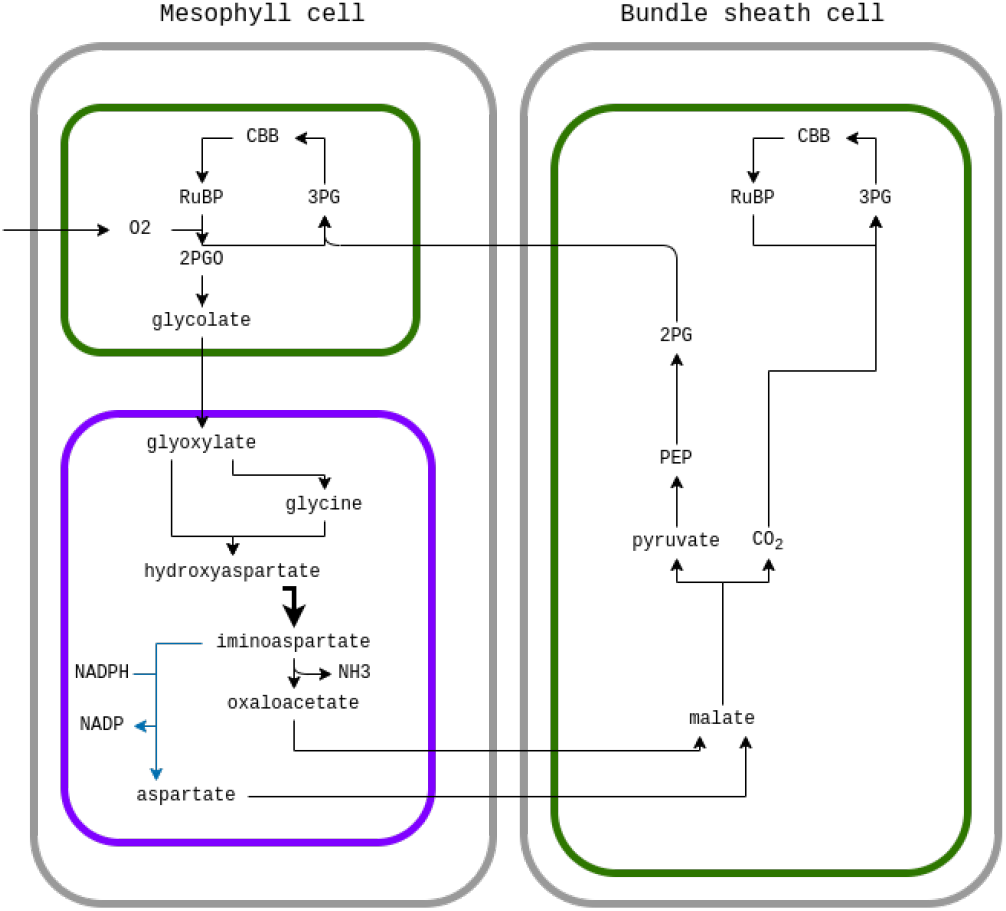
Schematic drawing of the BHAC pathway.

In order to decide on the reaction direction and whether we considered the reactions to be irreversible or reversible, we first calculated the equilibrium constants using the eQuilibrator tool, shown in table S28 [53].

**Table S28:** Standard Gibbs free energy of reaction and equilibrium constants for reactions in the BHAC pathway. These values were calculated using the eQuilibrator tool, with an assumed pH of 7.5, pMg of 3.0 and ionic strength of 0.25 M.

| Reaction | $\Delta_r G'^o$ | $K_{eq}$ |
| --- | --- | --- |
| hydroxyaspartate aldolase | -3.6 | 4 |
| hydroxyaspartate hydrolase | -3.5 | 4 |
| spontaneous reaction | -22.0 | 7.1e3 |
| aspartate oxidoreductase | -58.2 | 1.6e10 |
| aspartate aminotransferase | 2.8 | 0.3 |
| malate dehydrogenase | -26.5 | 4.4e4 |
| malic enzyme | 4.1 | 0.2 |
| pyruvate phosphate dikinase | 11.5 | 9.6e-3 |
| enolase | 3.8 | 0.2 |
| phosphoglycerate mutase | -4.5 | 6 |

The only reaction that was deemed irreversible was aspartate oxidoreductase, all other enzymatic reactions were modelled using reversible Michaelis-Menten kinetics, see section S3.1.1.

The implementation of the BHAC pathway requires addition of the pathway compounds hydroxyaspartate, iminoaspartate, oxaloacetate, aspartate, malate, pyruvate, PEP, as well as HCO3. The pathway compounds and HCO3 were implemented as variables. The additional parameters not present in the base model, can be found in table S30.

As HCO3 was implemented as a dynamic variable, it was necessary to add a carbonic anhydrase reaction to the model, for which we used diffusion kinetics kf_hydration 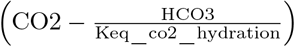.

#### S3.1.1 BHAC pathway model changes

##### Variables

**Table S29:** Variables of BHAC pathway model

| Model name | Name |
| --- | --- |
| HCO3 |  |
| asp | aspartate |
| glc2p | 2-phospho-D-glycerate |
| hasp | hydroxyaspartate |
| iasp | iminoaspartate |
| mal | malate |
| oxac | oxaloacetate |
| pep | phosphoenolpyruvate |
| pyr | pyruvate |

##### Parameters

**Table S30:** Parameters of BHAC pathway model

| Parameter name | Parameter value |
| --- | --- |
| AMP | 3.00e-01 |
| E0_aspat | 1.00e+00 |
| E0_aspor | 1.00e+00 |
| E0_enolase | 1.00e+00 |
| E0_gdc | 0.00e+00 |
| E0_haa | 1.00e+00 |
| E0_hah | 1.00e+00 |
| E0_mdh | 1.00e+00 |
| E0_me | 1.00e+00 |
| E0_oxaloacetate_formation | 1.00e+00 |
| E0_pgm | 1.00e+00 |
| E0_ppdk | 1.00e+00 |
| k_carbonic_anhydrase | 1.00e+03 |
| kcat_aspat | 8.40e+01 |
| kcat_aspor | 1.00e+00 |
| kcat_enolase | 1.00e+00 |
| kcat_haa | 1.00e+00 |
| kcat_hah | 1.00e+00 |
| kcat_mdh | 1.00e+00 |
| kcat_me | 3.90e+01 |
| kcat_oxaloacetate_formation | 1.00e+00 |
| kcat_pgm | 1.00e+00 |
| kcat_ppdk | 1.00e+00 |
| keq_aspat | 3.00e-01 |
| keq_ca | 5.00e+01 |
| keq_enolase | 2.00e-01 |
| keq_haa | 4.00e+00 |
| keq_hah | 4.00e+00 |
| keq_mdh | 4.40e+04 |
| keq_me | 2.00e-01 |
| keq_oxaloacetate_formation | 7.10e+03 |
| keq_pgm | 6.00e+00 |
| keq_ppdk | 9.60e-03 |
| km_aspat_p | 3.88e+00 |
| km_aspat_s | 2.53e+00 |
| km_aspor_s | 1.00e+00 |
| km_enolase_p | 1.00e+00 |
| km_enolase_s | 1.00e+00 |
| km_haa_p | 2.30e+00 |
| km_haa_s | 1.00e+00 |
| km_hah_p | 1.00e+00 |
| km_hah_s | 1.00e+00 |
| km_mdh_p | 1.00e+00 |
| km_mdh_s | 1.00e+00 |
| km_me_p | 1.25e-03 |
| km_me_s | 3.00e-03 |
| km_oxaloacetate_formation_p | 1.00e+00 |
| km_oxaloacetate_formation_s | 1.00e+00 |
| km_pgm_p | 1.00e+00 |
| km_pgm_s | 1.00e+00 |
| km_ppdk_p | 1.00e+00 |
| km_ppdk_s | 1.00e+00 |
| ppi | 3.00e-01 |
| vmax_aspat | 8.40e+01 |
| vmax_aspor | 1.00e+00 |
| vmax_enolase | 1.00e+00 |
| vmax_haa | 1.00e+00 |
| vmax_hah | 1.00e+00 |
| vmax_mdh | 1.00e+00 |
| vmax_me | 3.90e+01 |
| vmax_oxaloacetate_formation | 1.00e+00 |
| vmax_pgm | 1.00e+00 |
| vmax_ppdk | 1.00e+00 |

##### Derived Parameters

**Table S31:** Derived parameters of BHAC pathway model

| Name | Equation |
| --- | --- |
| vmax_aspat | $kcat\_aspat \cdot E0\_aspat$ |
| vmax_aspor | $kcat\_aspor \cdot E0\_aspor$ |
| vmax_enolase | $kcat\_enolase \cdot E0\_enolase$ |
| vmax_haa | $kcat\_haa \cdot E0\_haa$ |
| vmax_hah | $kcat\_hah \cdot E0\_hah$ |
| vmax_mdh | $kcat\_mdh \cdot E0\_mdh$ |
| vmax_me | $kcat\_me \cdot E0\_me$ |
| vmax_oxaloacetate_formation | $kcat\_oxaloacetate\_formation \cdot E0\_oxaloacetate\_formation$ |
| vmax_pgm | $kcat\_pgm \cdot E0\_pgm$ |
| vmax_ppdk | $kcat\_ppdk \cdot E0\_ppdk$ |

##### Reactions

###### aspartate aminotransferase (ASPAT)

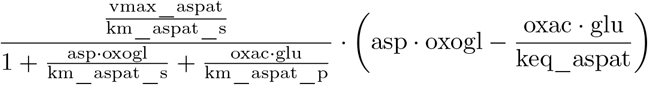

###### aspartate oxidoreductase (ASPOR)

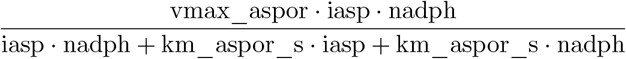

###### carbonic anhydrase (CA)

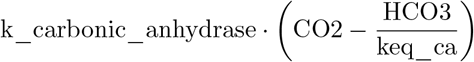

###### enolase (ENOLASE)

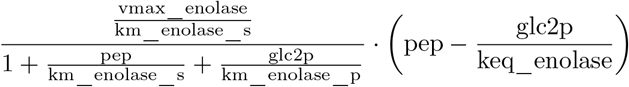

###### hydroxyaspartate aldolase (HAA)

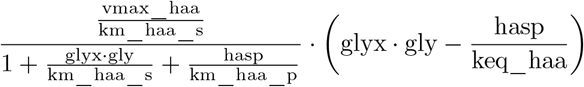

###### hydroxyaspartate hydrolase (HAH)

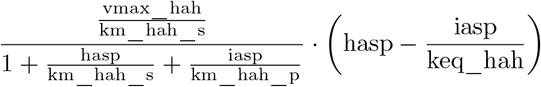

###### malate dehydrogenase (MDH)

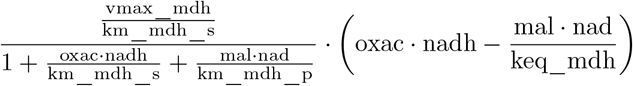

###### malic enzyme (ME)

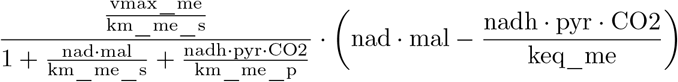

###### spontaneous oxaloacetate formation (OAF)

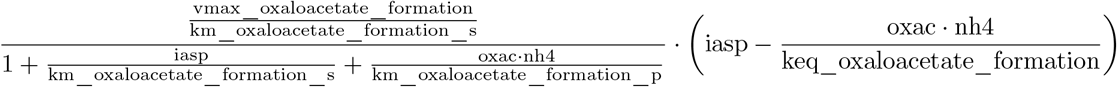

###### phosphoglucomutase (PGM)

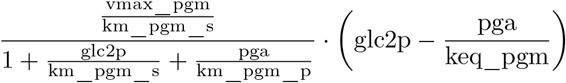

###### pyruvate-phosphate dikinase (PPDK)

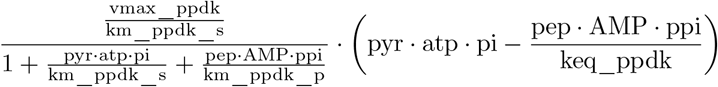

###### Stoichiometries

**Table S32:**
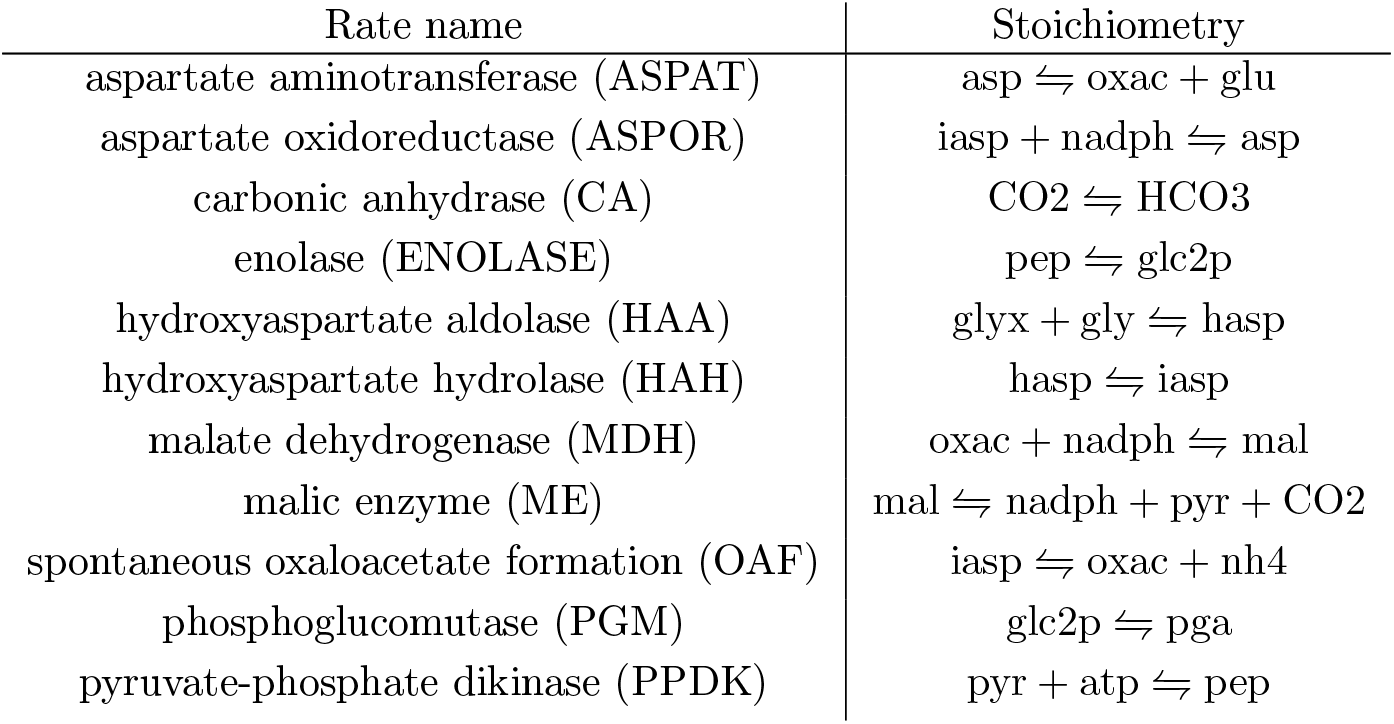
Stoichiometries of BHAC pathway model

## S4 Analysis of BHAC cycle

In order to critically assess the BHAC alternative pathway we performed systematic scans over a wide range of atmospheric carbon dioxide concentrations as well as illumination. This is important, as environmental conditions on Earth can vary quite dramatically both due to geographical considerations as well as climate change. Alternative pathways need to perform well in all of these conditions in order to be future-proof and commercially viable.

Fig. S16a shows a systematic parameter scan of atmospheric carbon dioxide concentration between 80 and 140 % of the reference condition (400 ppm) at a typical illumination of 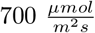. The BHAC pathway outperforms wildtype photorespiration in all of these conditions. However, the performance gain over the wildtype is most pronunced in low CO2 conditions and decreases with higher CO2 concentrations. This is to be expected, as with open stomata a higher external CO2 concentration will yield to a higher interal CO2 concentration (C_*i*_) which in turn will lead to less photorespiration overall.

**Figure S16:**
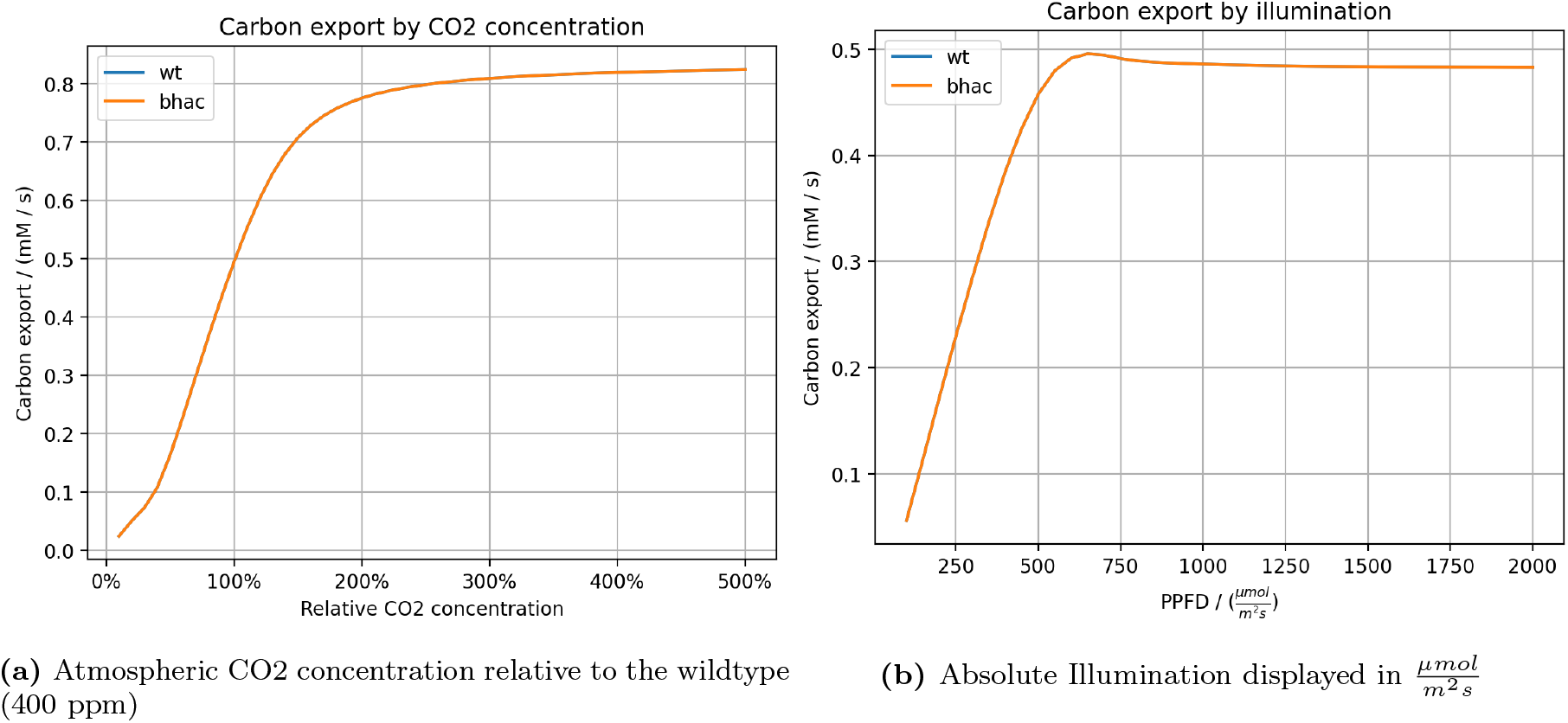
Comparison of the effect of atmospheric CO2 concentration (left) and illumination (right) on carbon export between the photorespiratory wildtype and the BHAC pathway. Carbon export is displayed in absolute units of 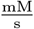.

Fig. S16b shows a systematic parameter scan of illumination between 100 and 2000 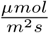 at a typical external carbon dioxide concentration of 400 ppm. In low light conditions the BHAC alternative pathway leads to the same carbon export as wildtype photorespiration. However, around an illumination of 400 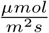, the carbon export of BHAC exceeds that of wildtype photorespiration. These results suggest that the BHAC alternative pathway is best suited for moderate to high-light conditions and may increase both yield under high light, as well as the maximal productive illumination. This is beneficial both for growing plants in regions with strong natural illumination as well as greenhouse plants, which can use additional light to grow faster - albeit at higher energy consumption.

Fig. S17a and Fig. S17b show metabolic control analyses of the BHAC alternative pathway over a wide range of illumination on the rate of rubisco carboxylation and carbon export respectively. Specifically, the normalised flux control coefficients of the enzymes with the highest absolute values are displayed. Those coefficients describe the amount of change to the flux (rate) of an enzyme that is to be expected when a parameter of another reaction (e.g. the enzyme concentration of the respective reaction) is changed. As there are a lot of parameters in the model, we only display coefficients which have an absolute value larger than 1 *×* 10^−2^.

The control on the rubisco flux is negligible for all other enzyme parameters in the model, as can bee seen in Fig. S17a, with the largest normalised flux control coefficients indicating that a 1 % increase in the enzyme concentration would only lead to a 0.06 % increase in the flux of the reaction. This intuitively makes sense, as plants can be assumed to have optimised the CBB cycle in a way to use all available carbon dioxide if no other limitations like drought are present.

Interestingly, while the rubisco flux itself is insensitive to changes in any enzyme concentrations, the carbon export can be very sensitive to changes in enzyme concentrations (see Fig. S17b). While increasing fructose-1,6-bisphosphatase (FBPase) by 1 % is predicted to increase carbon export by up to 6 % in certain conditions, increasing SBPase by 1 % will roughly decrease the carbon export to 10 % A possible explanation for this would be that with the additional supply of 3PGA by the BHAC alternative pathway, SBPase starts to have a high control of the CBB cycle (e.g. becomes a bottleneck reaction), which leads to an accumulation of triose phosphates or hexoses, which can then be exported and used by the plant.

**Figure S17:**
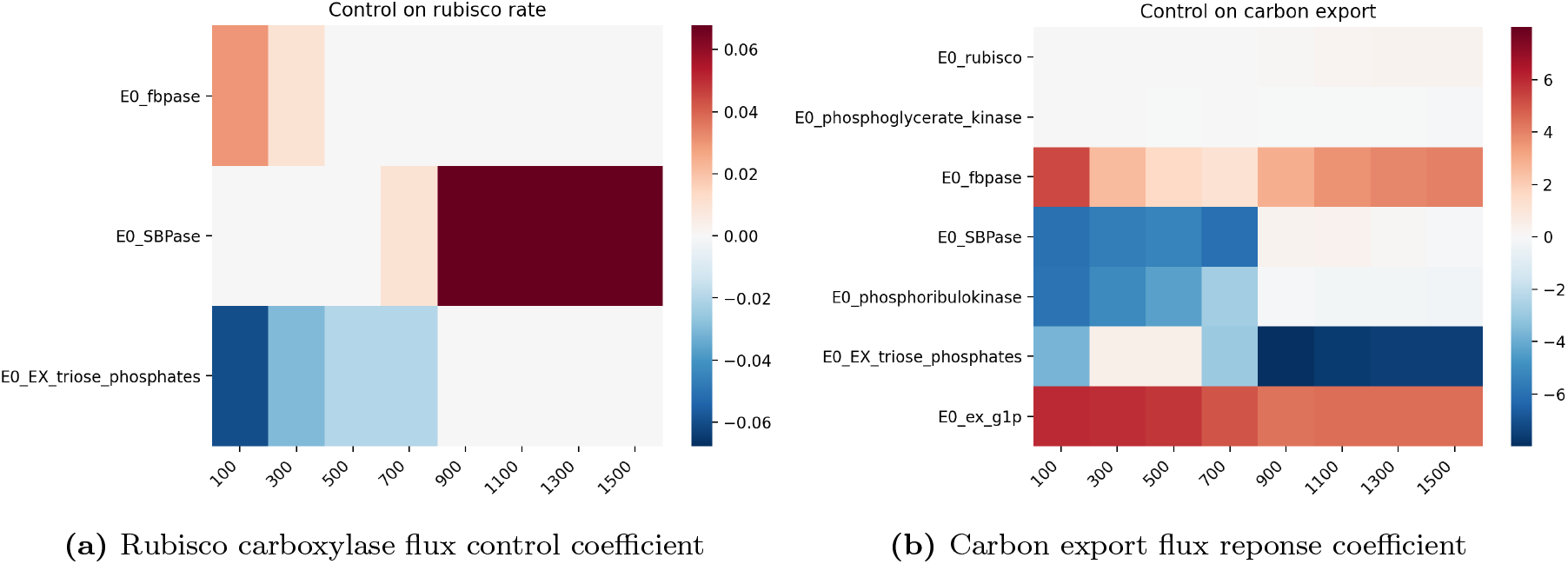
Normalised flux control and response coefficients of selected enzymes over multiple different light conditions between 100 and 1500 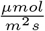. Red colors indicate positive control, while blue colors indicate negative control, with white colors representing no control of the respective flux whatsoever. Enzymes were selected by filtering out all coefficients below an absolute value below 1 *×* 10^−2^.

### TaCo model

The taco pathway pathway required addition of four reactions to the model, namely glycolyl-coa synthetase, glycolyl-coa carboxylase, tartronyl-coa reductase and tartronate-semialdehyde reductase [13, 14]. See Fig. S18 for an illustration.

**Figure S18:**
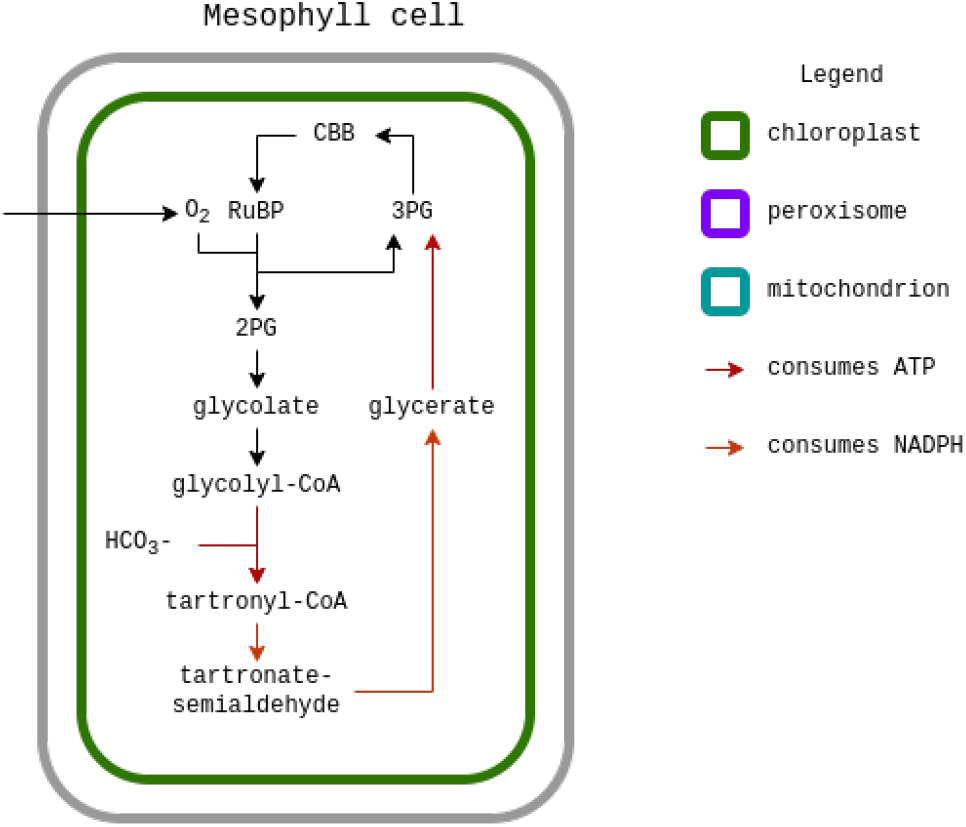
Schematic drawing of the TaCo pathway

These reactions have the following stoichiometries

**Table S33:** Stoichiometries of TaCo reactions

| Reaction | Stoichiometry |
| --- | --- |
| glycolyl-coa synthetase | $\text{ATP} + \text{CoA} + \text{glycolate} \leftarrow \text{Diphosphate} + \text{AMP} + \text{glycolyl-CoA}$ |
| glycolyl-coa carboxylase | $\text{glycolyl-CoA} + \text{ATP} + \text{HCO}_3^- \leftarrow \text{tartronyl-CoA} + \text{ADP} + \text{P}_i$ |
| tartronyl-coa reductase | $\text{NADPH} + \text{tartronyl-CoA} \leftarrow \text{NADP} + \text{CoA} + \text{tartronate-semialdehyde}$ |
| tartronate-semialdehyde reductase | $\text{tartronate-semialdehyde} + \text{NADH} \leftarrow \text{Glycerate} + \text{NAD}$ |

Note however, that GCC can use additional ATP in a futile ATP hydrolysis. We thus differentiated three cases: the optimal case, in which no futile ATP hydrolysis occurs, as well as the engineered M5 and L100N GCC variants, which use 3.9 and 1.7 ATP per reaction respectively. There is a tradeoff for the reduced futile ATP hydrolysis of the L100N variant to the increased catalytic constant though: while the M5 variants has a catalytic constant of 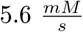, the L100N variants only features a catalytic constant of 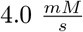.

In order to decide on the reaction direction and whether we considered the reactions to be irreversible or reversible, we first calculated the equilibrium constants using the eQuilibrator tool [53], shown in table S34.

**Table S34:** Standard Gibbs free energy of reaction and equilibrium constants for reactions in the TaCo pathway. These values were calculated using the eQuilibrator tool, with an assumed pH of 7.5, pMg of 3.0 and ionic strength of 0.25 M.

| Reaction | $\Delta_r G'^{\circ}$ | $K_{eq}$ |
| --- | --- | --- |
| glycolyl-coa synthetase | 9.25 | 0.024 |
| glycolyl-coa carboxylase | -20.86 | 4515.62 |
| tartronyl-coa reductase | 29.78 | $6.06e-6 \frac{1}{mM}$ |
| tartronate-semialdehyde reductase | -29.8 | 1.6e5 |

Only GCC and tartronate-semialdehyde reductase (TSR) have a negative ∆_*r*_G^*′*°^, however both glycolyl-CoA synthetase (GCS) and TSR can run forwards, given appropriate, but biologically feasible, concentrations of the substrates and products.

As neither of these equilibrium constants was deemed large or small enough to assume an irreversible reaction, we decided to model all of the reactions using reversible Michaelis-Menten kinetics, see section S4.1.1 for details. The pathway requires addition of the three pathway compounds glycolyl-CoA, tartronyl-CoA and tartronate-semialdehyde as well as the four additional ones AMP, diphosphate (PPi), HCO3 and CoA. The pathway compounds and HCO3 were implemented as variables, while AMP and PPi and CoA were all implemented as constants; their values, along with the other parameters not present in the base model, can be found in table S36. [53] As HCO3 was implemented as a dynamic variable, it was necessary to add a carbonic anhydrase reaction to the model, for which we used diffusion kinetics.

#### S4.1.1 TaCo pathway model changes

##### Variables

**Table S35:** Variables of TaCo pathway model

| Model name | Name |
| --- | --- |
| HCO3 |  |
| glyc-coa | glycolyl-CoA |
| tcoa | tartronate-CoA |
| tsa | tartronate-semialdehyde |

##### Parameters

**Table S36:**
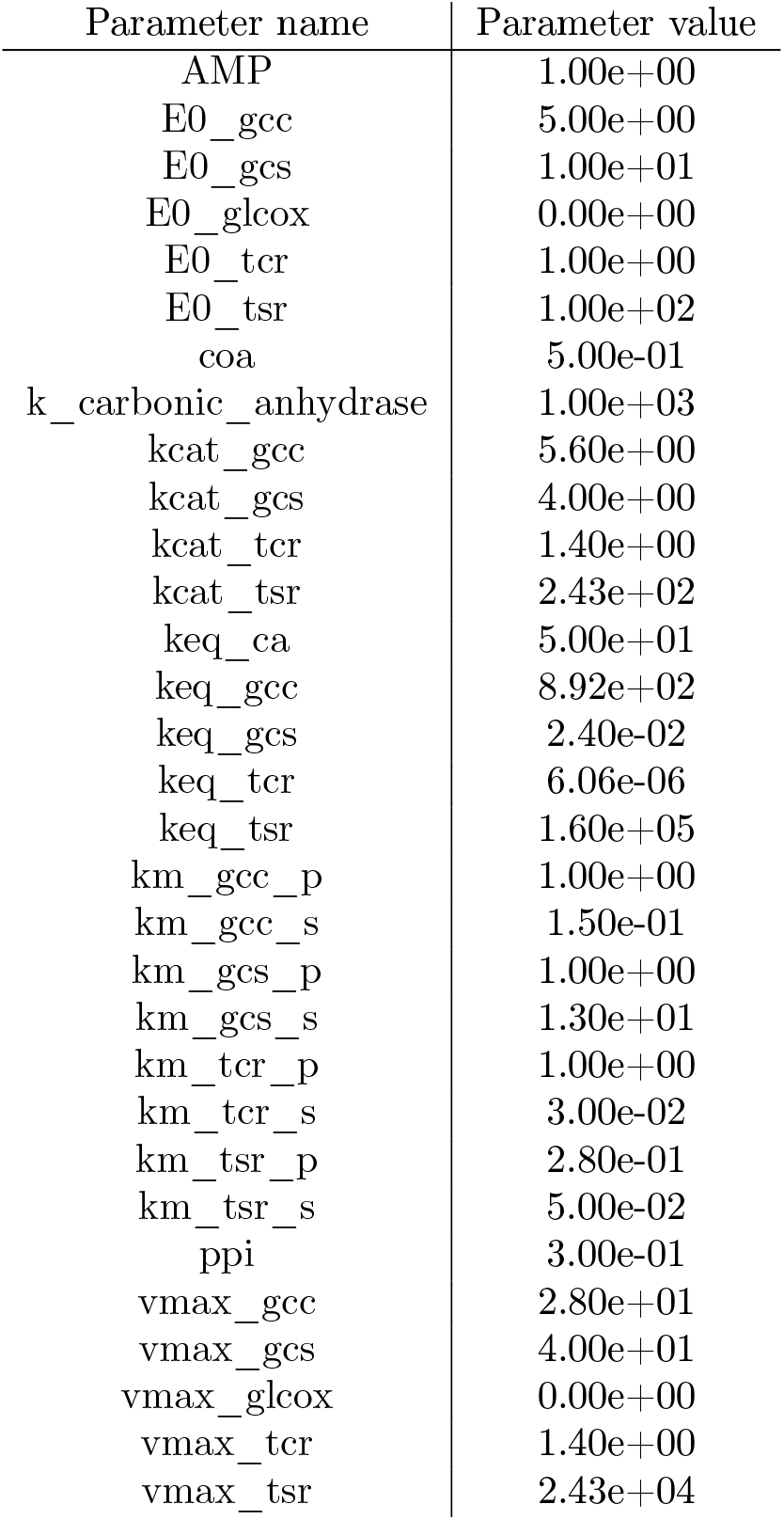
Parameters of TaCo pathway model

| Parameter name | Parameter value |
| --- | --- |
| AMP | 1.00e+00 |
| E0_gcc | 5.00e+00 |
| E0_gcs | 1.00e+01 |
| E0_glc ox | 0.00e+00 |
| E0_tcr | 1.00e+00 |
| E0_tsr | 1.00e+02 |
| coa | 5.00e-01 |
| k_carbonic_anhydrase | 1.00e+03 |
| kcat_gcc | 5.60e+00 |
| kcat_gcs | 4.00e+00 |
| kcat_tcr | 1.40e+00 |
| kcat_tsr | 2.43e+02 |
| keq_ca | 5.00e+01 |
| keq_gcc | 8.92e+02 |
| keq_gcs | 2.40e-02 |
| keq_tcr | 6.06e-06 |
| keq_tsr | 1.60e+05 |
| km_gcc_p | 1.00e+00 |
| km_gcc_s | 1.50e-01 |
| km_gcs_p | 1.00e+00 |
| km_gcs_s | 1.30e+01 |
| km_tcr_p | 1.00e+00 |
| km_tcr_s | 3.00e-02 |
| km_tsr_p | 2.80e-01 |
| km_tsr_s | 5.00e-02 |
| ppi | 3.00e-01 |
| vmax_gcc | 2.80e+01 |
| vmax_gcs | 4.00e+01 |
| vmax_glc ox | 0.00e+00 |
| vmax_tcr | 1.40e+00 |
| vmax_tsr | 2.43e+04 |

##### Derived Parameters

**Table S37:**
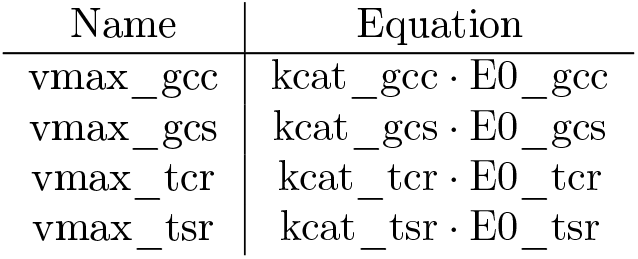
Derived parameters of TaCo pathway model

| Name | Equation |
| --- | --- |
| vmax_gcc | $kcat\_gcc \cdot E0\_gcc$ |
| vmax_gcs | $kcat\_gcs \cdot E0\_gcs$ |
| vmax_tcr | $kcat\_tcr \cdot E0\_tcr$ |
| vmax_tsr | $kcat\_tsr \cdot E0\_tsr$ |

##### Reactions

###### carbonic anhydrase (CA)

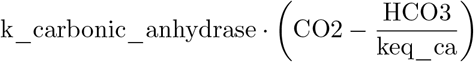

###### glycolyl-CoA carboxylase (GCC)

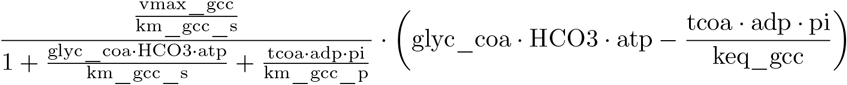

###### glycolyl-CoA synthetase (GCS)

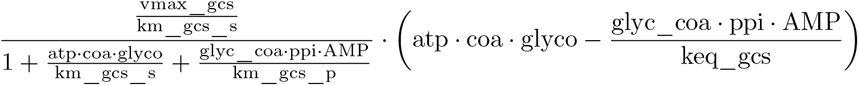

###### tartronyl-CoA reductase (TCR)

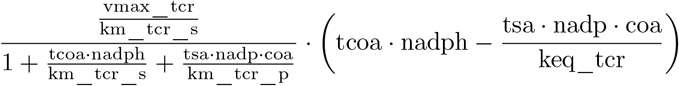

###### tartronate-semialdehyde reductase (TSR)

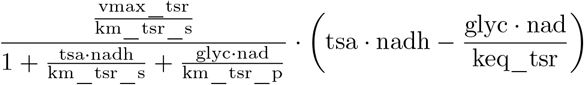

###### Stoichiometries

**Table S38:**
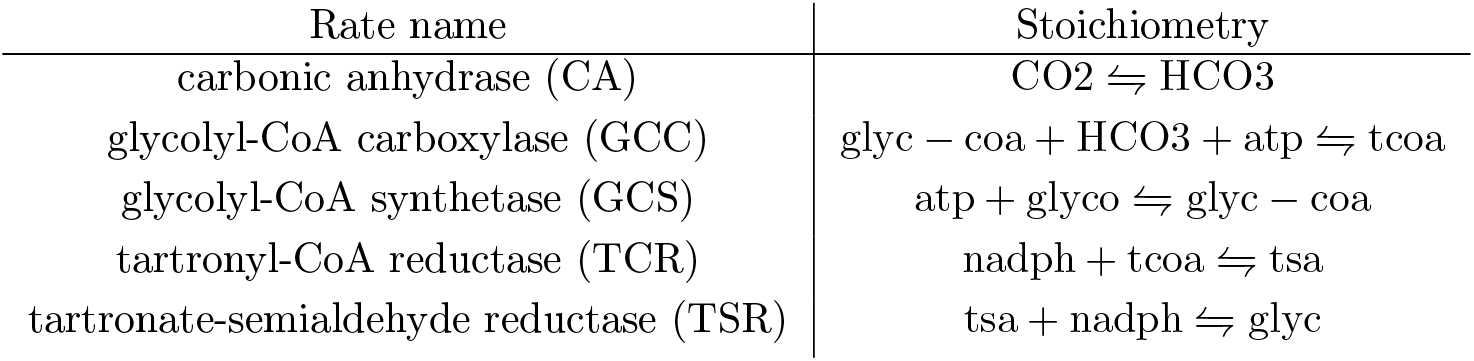
Stoichiometries of TaCo pathway model

